# Development and evaluation of PCR primers for environmental DNA (eDNA) metabarcoding of Amphibia

**DOI:** 10.1101/2021.10.29.466374

**Authors:** Masayuki K. Sakata, Mone U. Kawata, Atsushi Kurabayashi, Takaki Kurita, Masatoshi Nakamura, Tomoyasu Shirako, Ryosuke Kakehashi, Kanto Nishikawa, Mohamad Yazid Hossman, Takashi Nishijima, Junichi Kabamoto, Masaki Miya, Toshifumi Minamoto

## Abstract

Biodiversity monitoring is important for the conservation of natural ecosystems in general, but particularly for amphibians, whose populations are pronouncedly declining. However, amphibians’ ecological traits (e.g., nocturnal or aquatic) often prevent their precise monitoring. Environmental DNA (eDNA) metabarcoding—analysis of extra-organismal DNA released into the environment—allows the easy and effective monitoring of the biodiversity of aquatic organisms. Here, we developed and tested the utility of original PCR primer sets. First, we conducted *in vitro* PCR amplification tests with universal primer candidates using total DNA extracted from amphibian tissues. Five primer sets successfully amplified the target DNA fragments (partial 16S rRNA gene fragments of 160–311 bp) from all 16 taxa tested (from the three living amphibian orders Anura, Caudata, and Gymnophiona). Next, we investigated the taxonomic resolution retrieved using each primer set. The results revealed that the universal primer set “Amph16S” had the highest resolution among the tested sets. Finally, we applied Amph16S to actual metabarcoding and evaluated its detection capability by comparing the species detected using eDNA and physical survey (capture-based sampling and visual survey) in multiple agricultural ecosystems across Japan (160 sites in 10 areas). The eDNA metabarcoding with Amph16S detected twice as many species as the physical surveys (16 vs. 8 species, respectively), indicating the effectiveness of Amph16S in biodiversity monitoring and ecological research for amphibian communities.

## Introduction

Biodiversity loss has a major impact on the global environment (Butchart et al. 2010). This situation is particularly critical for freshwater organisms (Dudgeon et al. 2006) such as amphibians, which face serious threats, including habitat loss, water pollution, spread of infectious diseases, and the impact of invasive species (Stuart et al. 2004). As the decline in amphibian diversity is ongoing worldwide (Hof et al. 2011), their conservation is an urgent issue. Monitoring is important for collecting basic information (e.g., distribution) necessary for conservation (Costanza and Mageau 1999). Traditional amphibian monitoring includes physical surveys (capture or visual observation) and call surveys (for frogs; Heyer, Donnelly, Roy, Hayek, & Foster 2014); however, amphibian monitoring may be hindered by their small body size or ecological features such as cryptic or inaccessible habitat (e.g., muddy water, litter, subterrain, and forest canopy), opportunistic appearance (weather and seasonality), and nocturnal activity (Fellers et al. 2005).

Ficetola et al. (2008) first applied the environmental DNA (eDNA) technique to macro-organisms and succeeded in detecting the eDNA derived from the American bullfrog, *Lithobates catesbeianus*, from pond water. Subsequently, eDNA analysis was applied to the monitoring of various species and ecosystems (Goldberg et al. 2016, Deiner et al. 2017). As eDNA analysis allows for low-cost and extensive surveys (Jerde et al. 2011, Laramie et al. 2015), it has been used to monitor rare and endangered species (Bylemans et al. 2016, Sakata et al. 2017, Stat et al. 2018). Both species-specific eDNA analysis, which detects a specific species and estimates biomass (Takahara et al. 2012, Pilliod et al. 2013, Goldberg et al. 2018), and eDNA metabarcoding, which exhaustively detects target taxa by using high-throughput sequencing (HTS), have been applied for monitoring (Valentini et al. 2016, Yamamoto et al. 2017, Stat et al. 2018, Hayami et al. 2020). In eDNA metabarcoding for fish, universal primer sets have been sufficiently investigated and evaluated, and field applications are advanced (Miya et al. 2015, 2020, Deiner et al. 2017, Zhang et al. 2020, Miya 2021). For amphibia, several studies have used eDNA metabarcoding for biomonitoring various aquatic ecosystems, such as wetlands, tropical forests, and ponds (Valentini et al. 2016, Lopes et al. 2017, Sasso et al. 2017, Bálint et al. 2018, Kačergytė et al. 2021), or to investigate eDNA properties (Evans et al. 2016, Brys et al. 2021). Although several studies have used eDNA metabarcoding for amphibians, there has been no comparison or evaluation among universal primer sets as has been performed in eDNA metabarcoding studies on fish. For example, eDNA metabarcoding analysis using relatively short DNA sequences (<100 bp) occasionally provides genus-level resolution (Valentini et al. 2016). As amphibians have low migration capacity and occasionally have region-specific genetic characteristics (Nishizawa et al. 2011, Tominaga et al. 2013), developing a metabarcoding assay applicable to taxa with intraspecific genetic diversity is desirable. Therefore, the development of universal primer sets based on a proper evaluation of their characteristics is important.

In eDNA metabarcoding, the choice of PCR primers for amplification of target sequences (i.e., barcodes) is possibly one of the most influential factors in determining the detection probability of specific species or taxonomic groups (Freeland 2017, Alberdi et al. 2018). Ideally, universal PCR primers should have broad intra-group coverage, non-biased amplification across species, and high taxonomic resolution (Riaz et al. 2011). Some studies have evaluated the usability of the primers for fish eDNA metabarcoding (Bylemans et al. 2018, Collins et al. 2019, Zhang et al. 2020, Shu et al. 2021), focusing on several parameters such as amplification length, taxonomic coverage, primer universality, and interspecies resolution, which enables the use of appropriate primer sets that are consistent with the research objectives.

Here, we aimed to develop primer sets for amphibian metabarcoding assays with species- and/or subspecies-level resolution and to evaluate the effectiveness of the assay system herein proposed. To achieve these aims, we designed five universal primer sets for amphibians in the mitochondrial 16SrRNA region. We then evaluated them using several criteria, such as taxonomic resolution and taxonomic coverage, via in silico and in vitro tests. Finally, we examined the applicability and effectiveness of the best universal primer sets by conducting extensive field surveys and comparing the detected species between eDNA metabarcoding and physical surveys.

## Materials and Methods

### Primer development

#### Primer design

The 16S rRNA gene has been suggested as a suitable DNA barcoding marker for amphibians (Vences et al. 2005). To design universal primers for amphibian metabarcoding analyses, a total of 1,034 sequences of the mitochondrial 16S rRNA region of amphibians from 58 (/75: 77.3%) families and 293 (/564: 52.0%) genera was used from the National Center for Biotechnology Information database (NCBI, https://www.ncbi.nlm.nih.gov). To extract the taxonomic information of the 16S rRNA sequences from NCBI text (Genbank format), the program Taxonparser was used (newly written for this study; available from https://github.com/RyosukeKakehashi/taxonparser). The 16S rRNA sequences were aligned using MAFFT (Katoh and Standley 2013) with the default parameters of the MAFFT server (https://mafft.cbrc.jp/alignment/software/). Based on the alignments, several candidate universal primer sets were designed from the conserved regions (Fig. 1; Table 1), using Primer3web (Untergasser et al. 2012) and visual observation.

**Fig. 1.**
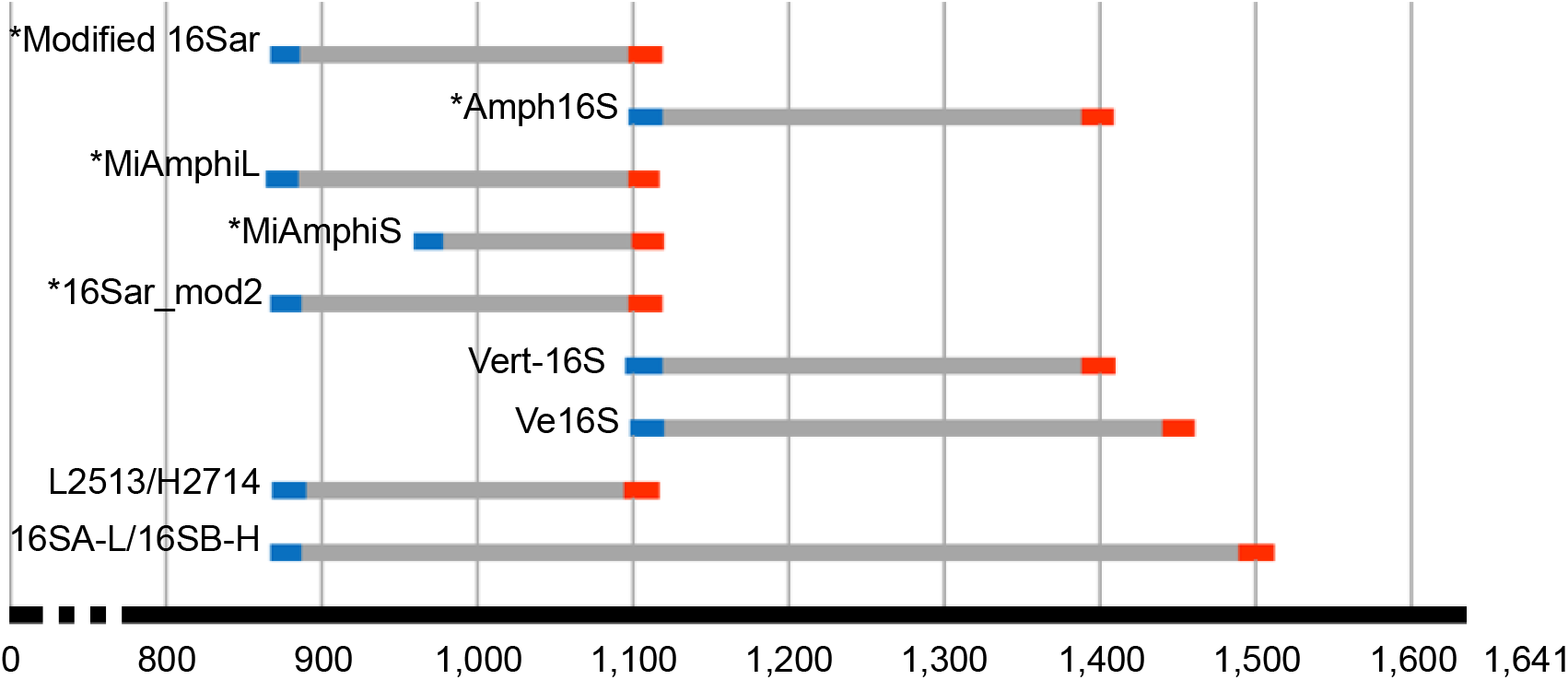
Locations of the nine amphibian metabarcoding primer pairs and amplicons on the target mitochondrial 16S rRNA gene. The target gene sequence of *Xenopus laevis* (Accession ID: MN259073.1) was used as template. Asterisk shows the primer set evaluated in this study. Note that the amplicon sizes of the primer sets may vary depending on the amphibian species. Others are existing primer sets: Vert-16S (Vences et al. 2016), Ve16S (Evans et al. 2016), L2513/H2714 (Kitano, Umetsu, Tian, & Osawa 2007), 16SA-L/16SB-H (Vences, Thomas, Van Der Meijden, Chiari, & Vieites 2005).

**Table 1.**
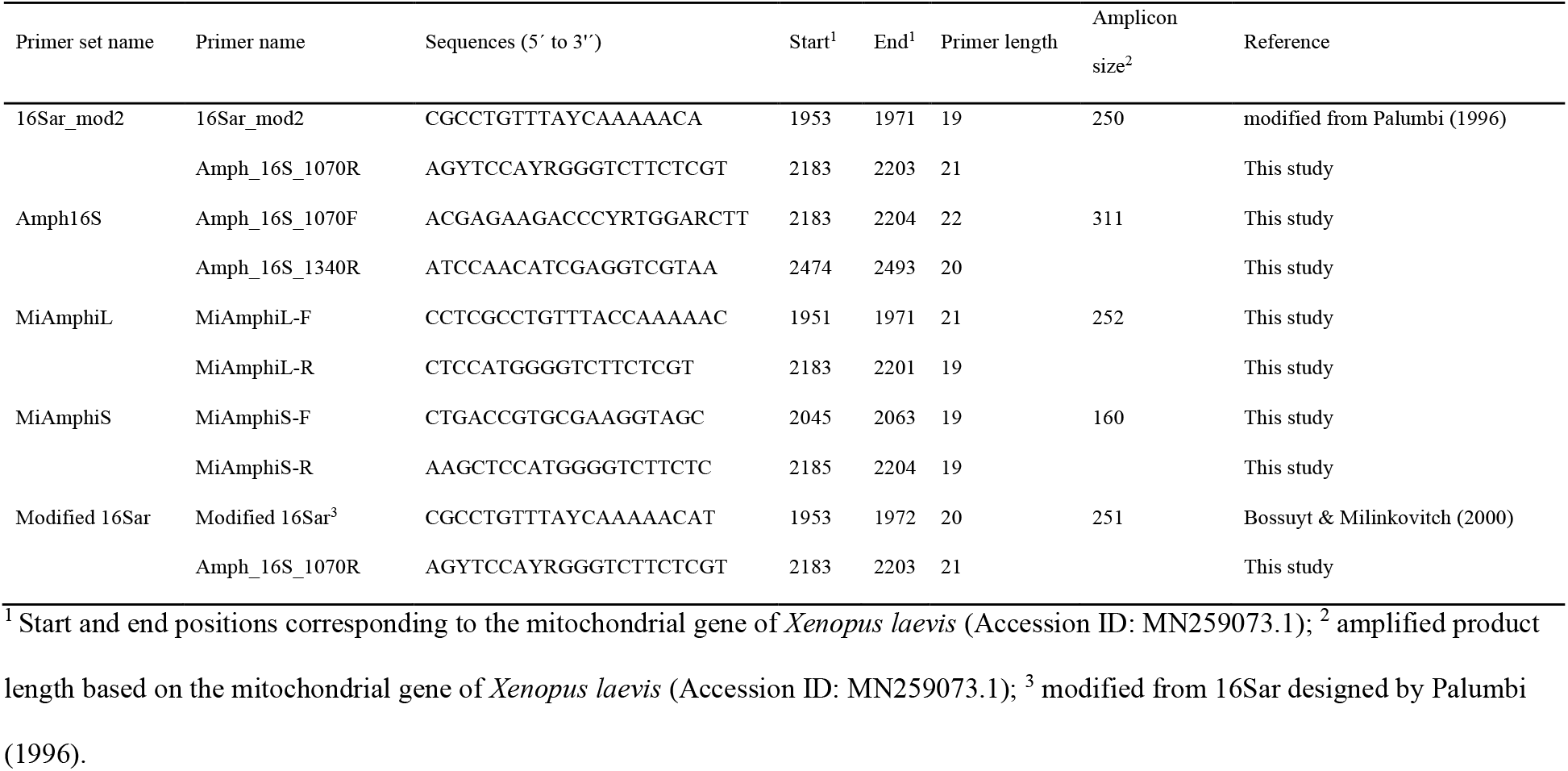
List of primers for the mitochondrial 16S rRNA gene region of amphibians evaluated in this study.

**Table 2.**
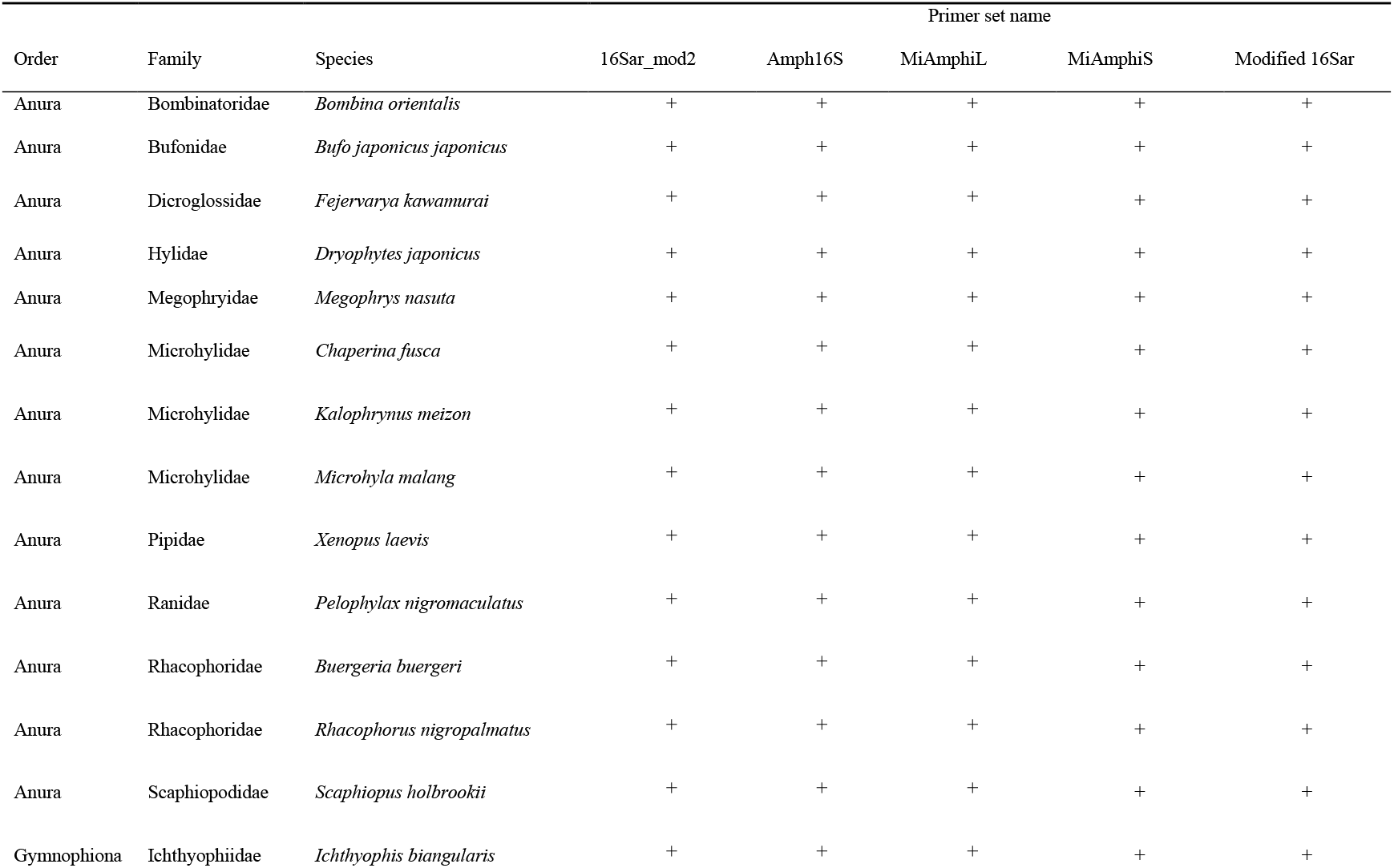
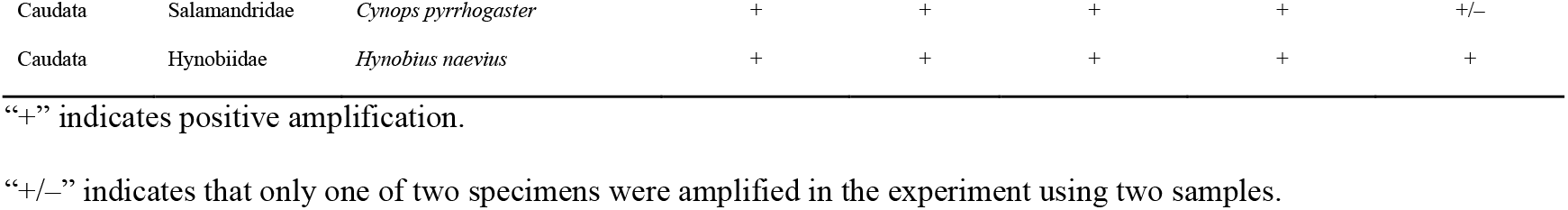
Results of the *in vitro* test for primer amplification.

#### *In silico* and *in vitro* evaluations of designed primer sets

To characterize each primer set, we performed *in silico* tests of the following three parameters: 1) universality of each priming site, 2) specificity of the priming site for target taxa, and 3) resolution of the internal amplified regions. To examine the universality of each primer, the frequency of bases at each locus of the primers was visualized using the sequence logo from the aforementioned 1,034 sequences. The specificity of each primer set was confirmed using *in silico* PCR, which was performed using the “search_pcr” command implemented in USEARCH v10.0.240 (Edgar 2010), targeting the full-length mitochondrial DNA reference database of vertebrates. This database was created from the mtDNA data deposited in NCBI (see Supporting Information) and comprises 26,096 sequences (fish: 7861, amphibia: 959, reptile: 1170, bird: 2988, mammalian: 13,118). For each primer set, *in silico* PCR was performed with minimum and maximum amplicon sizes of 100 and 1,000, respectively, and with varying maximum acceptable primer mismatches of 0, 1, and 2 (“maxdiff” option). The dissimilarity of the internal amplified regions of each primer set was calculated using the “dist.dna” function of the ape package in R ver. 3.6.3 (R Development Core Team 2019). The average length and dissimilarity of the internal amplified regions were then used to calculate the expected number of different bases among species in order to evaluate the taxonomic resolution of the primer sets. These numbers were compared among the primer sets by ANOVA and post-hoc Tukey-Kramer test.

*In vitro* tests were performed using tissue-derived DNA from 16 amphibians, including thirteen, one, and two species of Anura, Caudata, and Gymnophiona, respectively (Table S1). The total DNA used as PCR templates was extracted from previous studies and newly obtained in the present study (Kurabayashi et al. 2006, 2008, Kurabayashi and Sumida 2009, 2013). Five newly designed and existing primer sets were used (Table 1). The PCRs were conducted in a 10 μL solution containing 5 ng template DNA, 10x TaKaRa Ex Taq Buffer, 0.25 U TaKaRa Ex Taq DNA polymerase, 0.25 mM each dNTP, and 0.5 μM forward and reverse primers (each). PCR conditions were as follows: initial denaturation at 94°C for 30 s; 35 cycles of denaturation at 95°C for 30 s, annealing at 52°C for 30 s, and extension at 72°C for 30 s; and final extension at 70°C for 5 min. The resultant PCR solutions (5 μL) were electrophoresed on a 1.5% agarose gel to confirm amplification of the target DNA.

### Testing the effectiveness of universal primers

#### Field surveys

From July to September 2019, field surveys were conducted at 160 agricultural ecosystem sites across 10 areas of Japan (Fig. 2; Table S2). In each area, the study sites included various types of agricultural water environments, including streams, irrigation waterways, and ponds. These study areas covered a wide geographic region of Japan and included multiple local populations of some amphibian species. Most amphibians found in agricultural ecosystems are anurans, and these areas harbor approximately half of the Japanese anuran species (19/43 species) (Ministry of Agriculture, Forestry and Fisheries 2009; https://www.maff.go.jp/j/nousin/keityo/tanbo/). In 122 of the 160 sites, both eDNA and physical surveys (capturing and visual surveys) were conducted. For the remaining 38 sites, only the eDNA survey was performed.

**Fig. 2.**
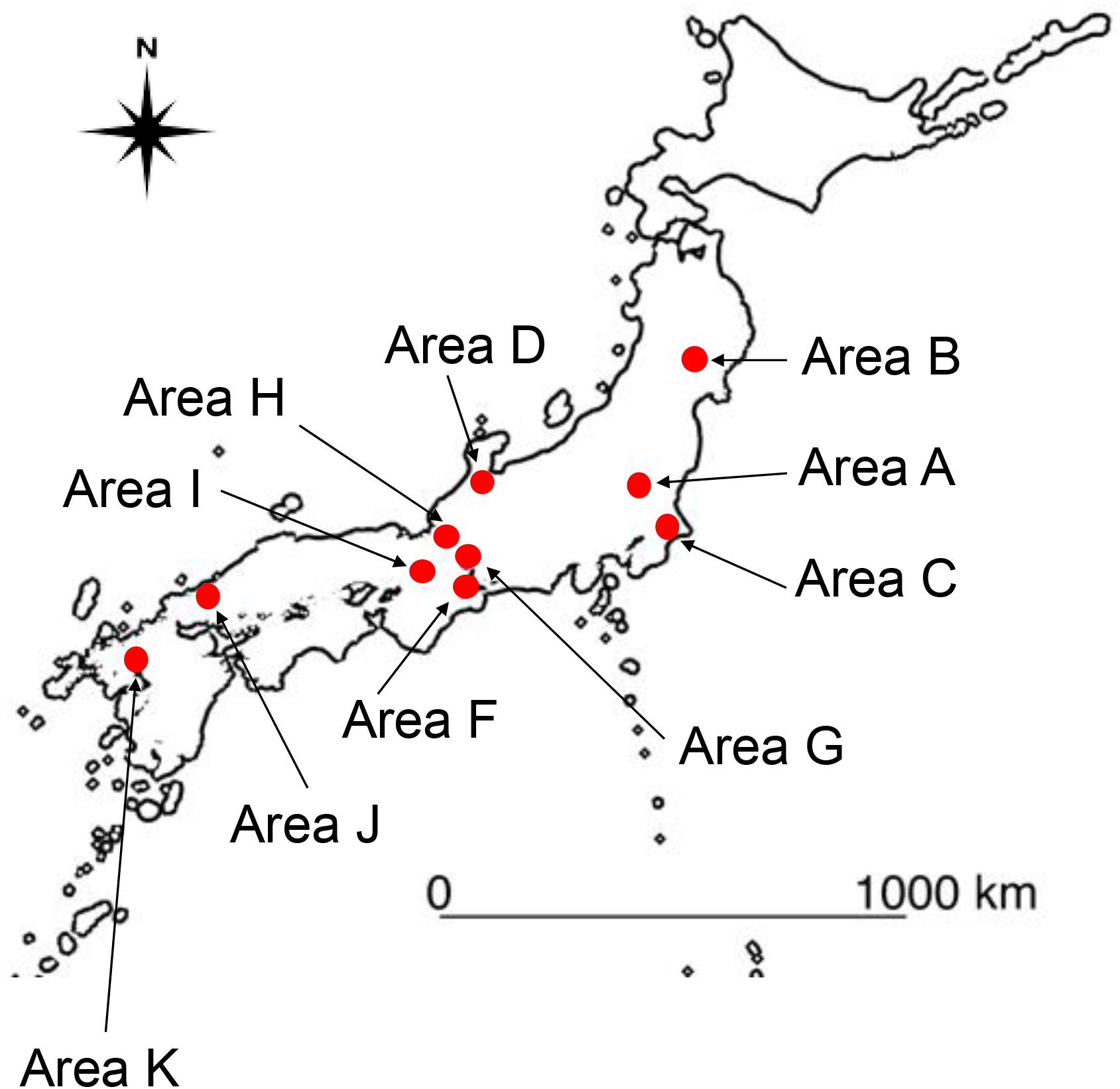
Map of the area surveyed in this study.

Water sampling was performed according to the Environmental DNA Sampling and Experiment Manual ver 2.1 (The eDNA Society, 2019). We sampled 1 L of surface water at each site, and after sampling, we added 1 mL benzalkonium chloride (final concentration = 0.1%) to prevent eDNA degradation (Yamanaka et al. 2017). To monitor potential contamination during the filtration and eDNA extraction processes, 1 L of distilled water was used as the negative control for each sampling date.

We performed physical surveys of amphibian species at the 122 sampling sites during the daytime (7:00–16:00) (Table S1) and compared the amphibian detectability between the physical and eDNA metabarcoding surveys. Physical surveys were conducted with constant capture efforts; specifically, for each site, three people collected amphibians for 30 min in a 50 m section. The collections were performed using a fixed net, hand net, net casting, and boxed net.

#### Environmental DNA sample processing

Filtration and eDNA extraction were performed according to the Environmental DNA Sampling and Experiment Manual ver 2.1 (The eDNA Society 2019). We filtered water samples using a glass fiber filter with nominal pore size of 0.7 μm (GF/F; GE Healthcare Life Science); however, a part of samples was filtered with two filters due to filter clogging. One or two filters were pooled into a single tube and preserved at −30°C. The eDNA on the filter was extracted using the Salivette (Sarstedt) and DNeasy Blood & Tissue Kit (QIAGEN Science, Hilden, Germany) and stored at −25°C. We obtained 100 μL of DNA eluted from the water sample. To prevent cross-contamination, all equipment used in the water collection and filtration steps, including plastic bottles, filter funnels, and tweezers, were decontaminated using >0.1% sodium hypochlorite solution (The eDNA Society 2019).

#### Application of the developed universal primers to environmental DNA analysis

To detect amphibian species from environmental samples, we amplified the partial 16SrRNA gene of amphibians and then performed high-throughput sequencing using a MiSeq platform (Illumina, San Diego, CA, USA). Among the primer sets, Amph16S— consisting of 16S rRNA gene specific primers (Amph16S_1070_F + Amph16S_1340_R)—was used to amplify the gene fragments from the eDNA samples for the following reasons: 1) this primer combination successfully amplified the target gene fragment of all 16 species tested belonging to all three amphibian orders (Table 3), and 2) the amplified region of this primer set was estimated to have the highest resolution in species identification among the primer sets tested (see Results). The first-round PCR (1st PCR) was performed in eight replicates with KOD-Plus-Neo polymerase (Toyobo, Osaka, Japan). Each PCR reaction (25 μL final volume) contained 300 nM primers, 2.5 μL 10x KOD buffer, 2.5 μL dNTPs (2 mM), 1.5 μL MgSO_4_ (25 mM), 0.5 μL KOD-Plus-Neo polymerase, 1 μL DNA template, and ultrapure water. The thermal cycle profile was 3 min at 95°C; 40 cycles of 20 s at 98°C, 15 s at 58°C, and 15 s at 72°C; and 72°C for 5 min. We used MiSeq-R1 tailed forward (5’-ACACTCTTTCCCTACACGACGCTCTTCCGATCTNNNNNN + gene-specific sequences (Amph16S_1070_F) – 3’ and R2 tailed reverse 5’-GTGACTGGAGTTCAGACGTGTGCTCTTCCGATCTNNNNNN + gene-specific sequences (Amph16S_1340_R) – 3’) primers. These primers included six random hexamers (N) to enhance cluster separation on the flow cells during initial base call calibrations on the MiSeq platform. Ultrapure water was used instead of eDNA in the eight reaction mixtures (non-template negative controls). After the eight technical replicates were pooled into a single tube, we removed unreacted reagents and primer dimers from the 1st PCR products using the GeneRead Size Selection Kit (QIAGEN Science, Hilden, Germany) according to the Environmental DNA Sampling and Experiment Manual (Version 2.1; The eDNA Society 2019).

**Table 3.** Detection (1) or not (0) of amphibian species by eDNA metabarcoding and physical surveys (phy).

| area | A |  | B |  | C |  | D |  | F |  | G |  | H |  | I |  | J |  | K |  |
| --- | --- | --- | --- | --- | --- | --- | --- | --- | --- | --- | --- | --- | --- | --- | --- | --- | --- | --- | --- | --- |
| Species/method | eDNA | phy | eDNA | phy | eDNA | phy | eDNA | phy | eDNA | phy | eDNA | phy | eDNA | phy | eDNA | phy | eDNA | phy | eDNA | phy |
| *Buergeria buergeri* | 0 | 0 | 0 | 0 | 0 | 0 | 0 | 0 | 0 | 0 | 0 | 0 | 0 | 0 | 0 | 0 | 0 | 0 | 1 | 0 |
| *Bufo japonicus* | 1 | 0 | 0 | 0 | 0 | 0 | 0 | 0 | 0 | 0 | 0 | 0 | 0 | 0 | 0 | 0 | 0 | 0 | 0 | 0 |
| *Bufo torrenticola* | 0 | 0 | 0 | 0 | 0 | 0 | 1 | 0 | 0 | 0 | 0 | 0 | 0 | 0 | 0 | 0 | 0 | 0 | 0 | 0 |
| *Fejervarya limnocharis* | 0 | 0 | 0 | 0 | 0 | 0 | 0 | 0 | 1 | 1 | 1 | 1 | 0 | 0 | 1 | 1 | 0 | 0 | 1 | 0 |
| *Dryophytes japonica* | 1 | 1 | 1 | 0 | 1 | 1 | 1 | 1 | 1 | 0 | 1 | 0 | 1 | 1 | 1 | 0 | 1 | 1 | 1 | 0 |
| *Lithobates catesbeianus* | 0 | 0 | 0 | 0 | 1 | 1 | 0 | 0 | 1 | 0 | 1 | 1 | 1 | 0 | 0 | 0 | 0 | 0 | 1 | 0 |
| *Pelophylax* sp. | 0 | 0 | 0 | 0 | 0 | 0 | 0 | 0 | 0 | 0 | 0 | 0 | 0 | 0 | 0 | 0 | 0 | 0 | 1 | 0 |
| *Pelophylax nigromaculatus* | 0 | 0 | 0 | 0 | 0 | 0 | 1 | 1 | 1 | 1 | 1 | 1 | 1 | 1 | 1 | 0 | 1 | 1 | 0 | 0 |
| *Pelophylax porosus* subsp. | 1 | 1 | 1 | 0 | 1 | 1 | 0 | 0 | 1 | 0 | 0 | 0 | 1 | 0 | 0 | 0 | 0 | 0 | 0 | 0 |
| *Rana japonica* | 1 | 1 | 0 | 0 | 1 | 0 | 0 | 0 | 0 | 0 | 0 | 0 | 0 | 0 | 0 | 0 | 1 | 1 | 1 | 0 |
| *Rana ornativentris* | 0 | 0 | 0 | 0 | 0 | 0 | 0 | 0 | 0 | 0 | 0 | 0 | 1 | 0 | 0 | 0 | 0 | 0 | 0 | 0 |
| *Rana tagoi tagoi* | 0 | 0 | 1 | 0 | 0 | 0 | 0 | 0 | 0 | 0 | 0 | 0 | 1 | 0 | 0 | 0 | 1 | 0 | 1 | 0 |
| *Zhangixalus arboreus* | 0 | 0 | 0 | 0 | 0 | 0 | 0 | 0 | 0 | 0 | 1 | 0 | 1 | 0 | 0 | 0 | 0 | 0 | 0 | 0 |
| *Zhangixalus schlegelii* | 0 | 0 | 1 | 0 | 1 | 0 | 0 | 0 | 0 | 0 | 0 | 0 | 0 | 0 | 0 | 0 | 1 | 0 | 1 | 0 |
| *Glandirana rugosa* | 1 | 1 | 1 | 1 | 0 | 0 | 1 | 1 | 1 | 0 | 0 | 0 | 1 | 0 | 1 | 0 | 1 | 1 | 1 | 0 |
| *Cynops pyrrhogaster* | 0 | 0 | 0 | 0 | 0 | 0 | 0 | 0 | 0 | 0 | 0 | 0 | 0 | 0 | 0 | 0 | 0 | 1 | 0 | 0 |

We carried out a second-round PCR (2nd PCR) using the purified products from the 1st PCR as templates. The 2nd PCR was performed using P5-i5-R1 (5’ – AATGATACGGCGACCACCGAGATCTACAXXXXXXXXACACTCTTTCCCTAC ACGACGCTCTTCCGATCT – 3’) and P7-i7-R2 (5’ – CAAGCAGAAGACGGCATACGAGATXXXXXXXXGTGACTGGAGTTCAGACG TGTGCTCTTCCGATCT – 3’) primers to add MiSeq adapter sequences and 8-bp index sequences to both ends of the amplicons. The octo-X segments represent dual-index sequences. The 2nd PCR was conducted in a 12 μL solution containing 6.0 μL of 2× KAPA HiFi HotStart ReadyMix (KAPA Biosystems, Wilmington, WA, USA), 3.6 pmol each primer, and 1 μL 1st PCR product. The PCR profile was as follows: 3 min at 95°C; 15 cycles of 20 s at 98°C, 15 s at 72°C, and 72°C for 5 min.

The 2nd PCR products were pooled in equal volumes into a single 1.5-ml tube. Target amplicons were selected by electrophoresis using E-Gel® SizeSelect 2% (Thermo Fisher Scientific, Waltham, MA, USA) with the E-Gel Precast Agarose Electrophoresis System (Thermo Fisher Scientific, Waltham, MA, USA). DNA concentration was measured using real-time PCR assays (QuantStudio3; Thermo Fisher Scientific, Waltham, MA, USA), and the library size distribution was confirmed using TapeStation 4200 (Agilent, Tokyo, Japan). The concentration of the DNA library was adjusted to 4 nM. Finally, the library was sequenced using an Illumina MiSeq v2 Reagent kit for 2× 250 bp PE (Illumina, San Diego, CA, USA).

#### Bioinformatics

Raw sequencing reads were converted to FASTQ format using Illumina bcl2fastq2 v.2.17 software allowing zero mismatches. To perform the species identification from the MiSeq output, FASTQ data were processed using the pipeline of the metabarcoding program package Claident version 0.2.2017.05.22 (Tanabe & Toju 2013; downloaded from https://www.claident.org/). We demultiplexed the data and removed any reads with low-quality index sequences (i.e., Phred score < 30) using the clsplitseq command with the option minqualtag = 30. Paired-end reads were then merged using the clconcatpair command with the option maxnmismatch = 20 and minovllen = 20. We then trimmed low-quality tails until the Phred scores of the last base were 30 or higher. We removed low-quality sequences (Phred score < 30) using the clfilterseq command with the options minqual = 30 and maxplowqual = 0.1. In addition, apparently noisy and chimeric reads were removed using the clcleanseqv command with the options primarymaxnmismatch = 0, secondarymaxnmismatch = 1, and pnoisycluster = 0.5. After quality control, we classified the reads with > 99% sequence similarity into a molecular operational taxonomic unit (MOTU) using the clclassseqv command with the option of minident = 0.99. We obtained denoising sequence data, and identified each MOTU as a species using online BLASTN (https://blast.ncbi.nlm.nih.gov/Blast.cgi) based on a 97% homology criterion.

MOTUs with less than 10 reads per sample were discarded because of potential contamination. The remaining sequence reads assigned to amphibians were vetted based on habitat, and species assignments were finalized. For all samples, the read depth was sufficiently large to saturate the number of amphibian species detected (checked using the “rarecurve” function in the vegan package version 2.5-4 (Oksanen et al. 2019)). Therefore, rarefaction was not performed.

#### Comparison with physical surveys

To compare the monitoring results between eDNA metabarcoding and physical surveys, the following analyses were performed using the vegan package version 2.5-4 and lme4 package version 1.1–21 (Bates et al. 2015) in R ver. 3.6.3 (R Development Core Team 2019). We used the VennDiagram package version 1.6.20 and ggplot2 package version 3.2.0.9000 to draw figures. The site-species read matrix was converted to the presence/absence of each species, and the data of 122 sites where both eDNA survey and physical surveys were carried out were used for comparison (Table S2). First, we compared the species composition of the monitoring methods. Non-metric multidimensional scaling (NMDS) with 10,000 permutations depicted the similarity of species composition among sites and methods measured using the Jaccard similarity coefficient, and PERMANOVA with 10,000 permutations was conducted using the “adonis” function in vegan. In addition, to compare the beta dispersion of the methods, PERMDISP analysis was performed with the “betadisper” function in vegan. Second, we compared the number of amphibian species detected between the monitoring methods. We fit a generalized linear mixed model with a Poisson distribution using the function “glmer.” In this model, the number of species was used as the response variable. The monitoring methods (i.e., eDNA metabarcoding or physical surveys) were set as explanatory variables, and the survey area ID was set as a random effect to consider differences in the number of species inhabiting the survey areas. Finally, we visualized the detected community composition between the two methods in each area by drawing a Venn diagram.

## Results

### Performance comparison of the PCR primer sets

We tested five sets of universal primer candidates for the amphibian 16S rRNA genes (Table 1). The sets of “MiAmphiS” and “Amphi16S” had the shortest and longest amplicon sizes, respectively. The universality of each primer set was shown by the sequence logo (Fig. S1). Both forward and reverse primers of “MiAmphiL” and “MiAmphiS” had no degenerated bases in both forward and reverse primers and high universality at the 3’-end of both primers; however, there were several mismatches in the region between the 5’-end and the center of both primers, which was located on a variable region among species. The primers of “Modified 16Sar” and “16Sar_mod2” contained some degenerated bases, resulting in high universality. Amph16S” had mixed bases in the forward primer and no mixed base in the reverse primer, and also showed high universality.

Regarding the power of amplification, we performed *in vitro* amplification tests for all five primer sets using DNA templates extracted from the 16 amphibian species belonging to all three amphibian orders, and all primer sets amplified the target fragments tested. We performed *in silico* PCR using vertebrate 16S rRNA gene data reported to date (7861 fishes, 959 amphibians, 1170 reptiles, 2988 birds, and 13,118 mammals). The specificity of each primer set examined by *in silico* PCR is shown in Table S3. When we did not allow any mismatch between the primers and target 16S rRNA gene sequences (maxdiff = 0), Amph16S showed the highest amplification rate (925/959 sequences; 96.4%). However, this primer set also had the highest amplification rate for non-amphibian taxa. MiAmphiL and MiAmphiS had the lowest amplification rates for non-amphibian taxa (3.1% and 5.4%, respectively), but they had the lowest amplification rates for amphibians. Even when we allowed two mismatches (maxdiff = 2), Amph16S showed the highest amplification rate for amphibians (98.7%). Under maxdiff = 2, the amplification rates for reptiles and birds were clearly lower for MiAmphiL and MiAmphiS (8.2% and 28.9% for reptiles and 0% and 4.9% for birds, respectively) than for Amph16S (90.3% and 99.4%, respectively), whereas the amplification rates for fishes and mammals did not differ significantly among these primer sets.

In the context of taxonomic resolution, Amph16S had the highest expected number of bases that differ among species (mean: 92.90, range: 68.77–163.23) (Fig. 3, Fig. S2), whereas MiAmphiS had the lowest value (mean: 23.00, range: 17.17–34.53). The means of 16S_mod2, MiAmphiL, and Modified 16Sar were 43.61, 43.50, and 44.89, respectively.

**Fig. 3.**
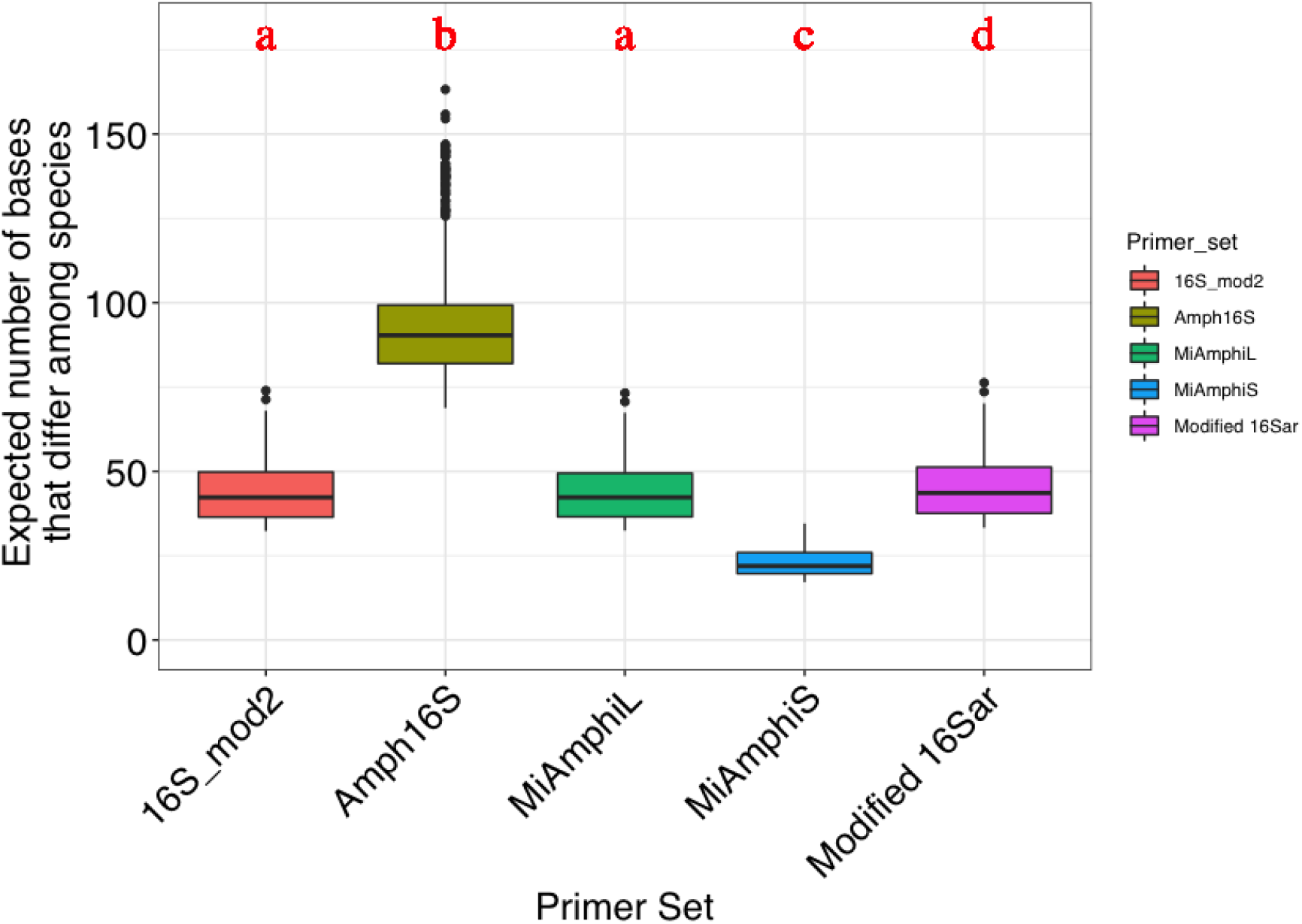
Comparison of the taxonomic resolutions among primer sets. The vertical axis indicates the expected number of bases that differ among species within the amplified region in each primer set. Each category indicates a set of primers; 16Sar_mod2: 16Sar_mod2 and Amph_16S_1070R, Amph16S: Amph_16S_1070F and Amph_16S_1340R, MiAmphiL: MiAmphiL-F and MiAmphiL-R, MiAmphiS: MiAmphiS-F and MiAmphiS-R, Modified 16Sar: Modified 16Sar and Amph_16S_1070R. The expected number of bases differed significantly among primer sets (ANOVA: *p* <0.05). Significant differences were indicated by different letters (Tukey-Kramer test: *p* < 0.05).

Amph16S had the highest universality for PCR amplification and taxonomic resolution in eDNA metabarcoding. Therefore, we regarded this primer set as the most useful and was applied in the subsequent field surveys.

### Evaluating the effectiveness of Amph16S in field survey

In the actual metabarcoding analysis using eDNA with Amph16S for the 160 field sites, we detected a total of 15 anuran and one caudate species from 122 water samples (see Discussion and Table S4). None of the negative controls detected any of the species. Eight amphibian species were detected in the physical surveys (Table S5). Combining the eDNA metabarcoding and physical surveys, 16 amphibian species were detected (Table 3). Among them, seven species were detected by both eDNA metabarcoding and physical surveys. Eight species (*Buergeria buergeri, Bufo japonicus, Bu. torrenticola, Pelophylax* sp., *Rana ornativentris, R. t. tagoi, Zhangixalus arboreus*, and *Z. schlegelii*) were detected only in the eDNA metabarcoding, and only one species (*Cynops pyrrhogaster)* was detected by physical surveys only. Seven sequences from multiple geographic areas were assigned to the single anuran species *Glandirana rugosa* (Fig. S3), although they showed a >3% nucleotide divergence. This is consistent with a previous report of many local populations with high genetic divergence in this species (Sekiya et al. 2013).

Community composition differed significantly between the eDNA metabarcoding and physical surveys (PERMANOVA: *p* <.001; Fig. 4; Table 4). The subsequent PERMDISP analysis further indicated that the amphibian community composition was significantly heterogeneous between the monitoring methods (*p* <.001; Table 5), and the number of species detected by eDNA metabarcoding was higher than that of physical surveys (GLMM: *p* <.001; Fig. 5A; Table 6). The amphibian community composition detected by the eDNA metabarcoding method encompassed that of the physical survey in all areas excluding area J (Fig. 5B; Table 3).

**Fig. 4.**
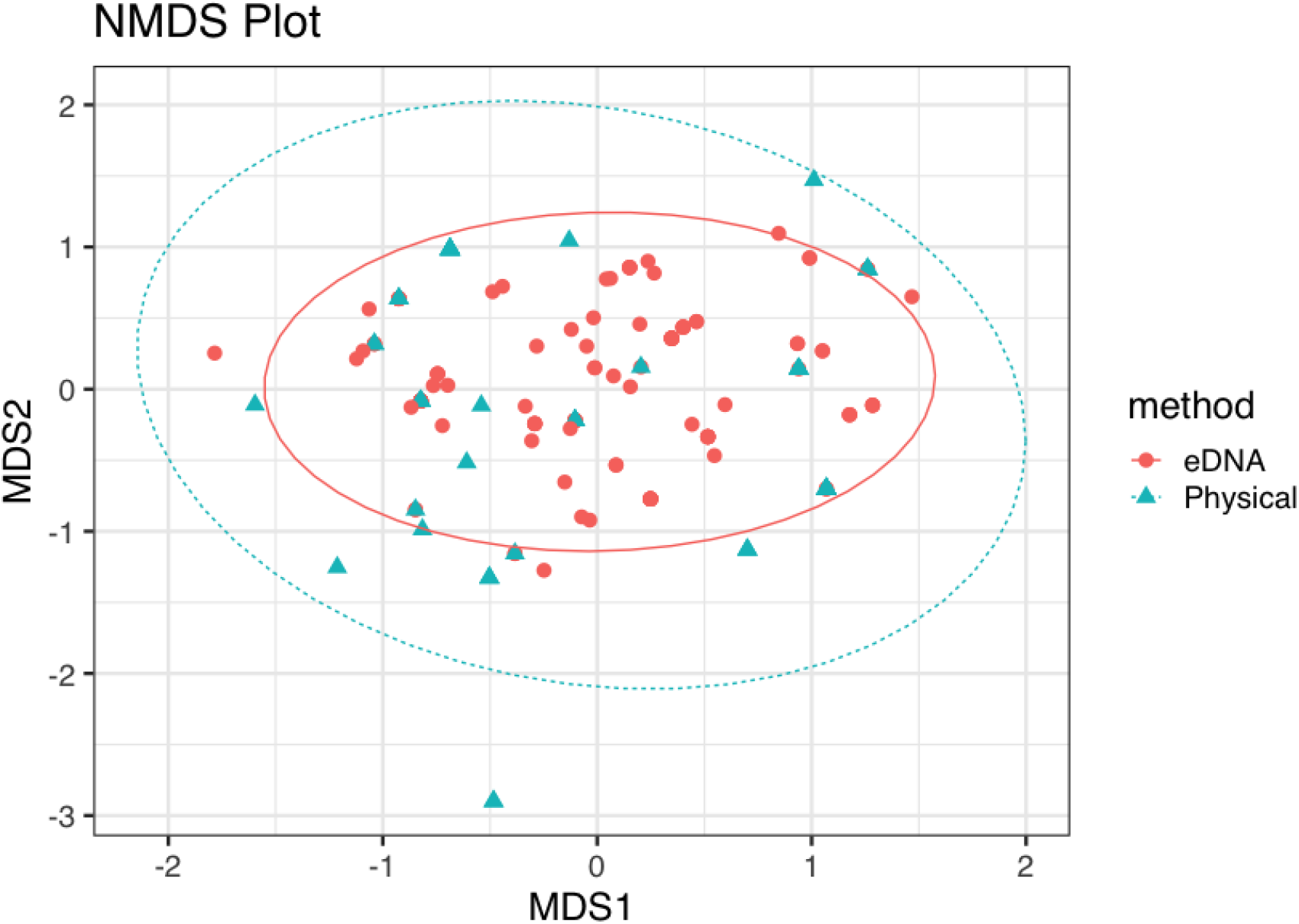
Comparison of amphibian community structures for each area and method. The NMDS plot showed variation of composition. The composition is significantly different between monitoring methods (PERMANOVA: *p* < 0.001). The ellipses show the 95% confidence level based on the centroid calculated for each monitoring method.

**Table 4.** Summary of the results of the PERMANOVA between monitoring methods (eDNA metabarcoding and physical survey) for amphibian community composition.

|  | df | Sums of Sqs | Mean Sqs | F | R<sup>2</sup> | *p* |
| --- | --- | --- | --- | --- | --- | --- |
| Methods | 1 | 1.746 | 1.74645 | 5.5987 | 0.03083 | <.001 |
| Residuals | 176 | 54.9 | 0.31194 |  | 0.96917 |  |
| Total | 177 | 56.65 |  |  |  |  |

**Table 5.** Summary of the results of the PERMDISP between monitoring methods (eDNA metabarcoding and physical survey) for amphibian community composition.

|  | df | Sums of Sqs | Mean Sqs | F | *p* |
| --- | --- | --- | --- | --- | --- |
| Methods | 1 | 0.1196 | 0.119603 | 12.024 | <.001 |
| Residuals | 176 | 1.7506 | 0.009947 |  |  |

**Fig. 5.**
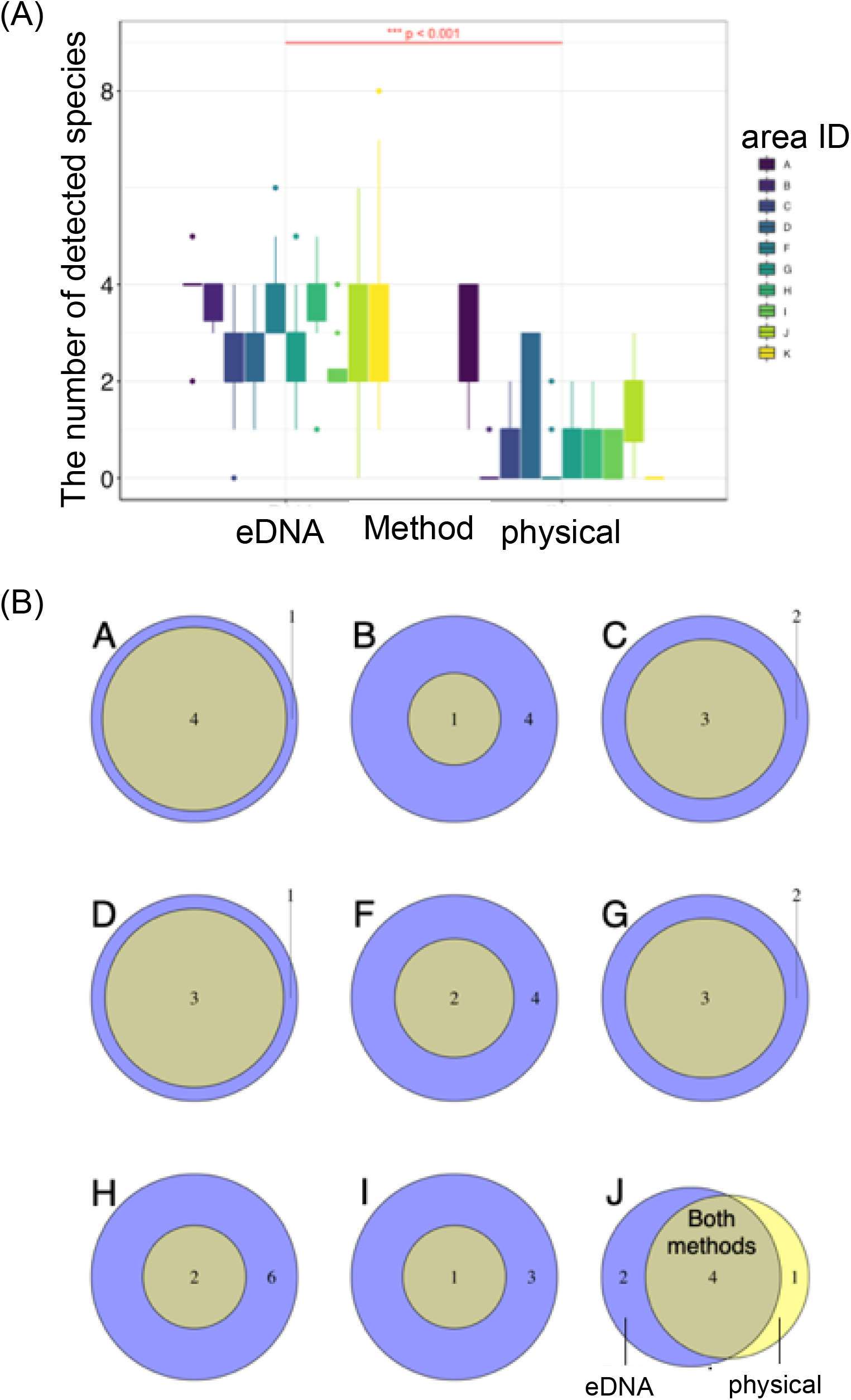
(A) Comparison of the number of species detected between monitoring methods. The number of species detected by eDNA metabarcodingis higher than one of physical surveys (GLMM: *p* < 0.001). (B) Comparison of community composition at each area via Venn diagrams. Blue and yellow show eDNA metabarcoding and physical surveys, respectively. The number indicates the number of detected species.

**Table 6.**
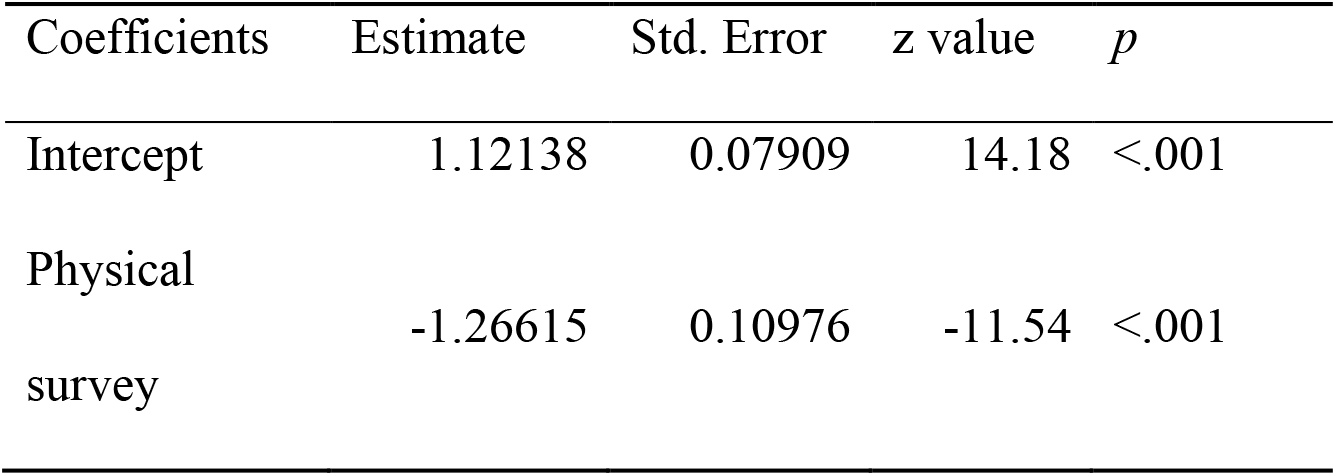
Result of the generalized mixed linear model for comparison of the number of species detected between monitoring methods.

## Discussion

With regard to the usability of universal primers for eDNA metabarcoding, amplicon length and consistency with the priming site are important factors. Long DNA fragments have a faster degradation rate than short DNA fragments (Jo et al. 2017), resulting in a disadvantage in detection sensitivity for the former. However, a long amplification length increases taxonomic resolution. Thus, there is a trade-off between detectability and resolution, depending on the amplification length. The introduction of degenerate bases is another point to consider in primer design. A primer with a degenerate site can reduce the mismatch between the primer and template DNA, but it may also cause non-specific amplification of non-target taxa (Zhang et al. 2020). Therefore, when designing and selecting primer sets, as herein performed, it is recommended to consider the sensitivity, taxonomic resolution, and specificity of each primer set through *in silico* and *in vitro* tests, and to select the primers according to the target ecosystem and purpose of the study.

The number of amphibian species detected using eDNA metabarcoding was higher than that from physical surveys. In addition, the amphibian community composition detected by eDNA metabarcoding encompassed that from physical surveys in almost all sampling areas. Similar to the results of previous studies on eDNA metabarcoding for fishes (Miya et al. 2015; Valentini et al. 2016; Yamamoto et al. 2017), metabarcoding had higher detectability of amphibian taxa than physical surveys. Compared to the current physical survey, only the red-bellied newt, *Cynopus pyrrhogaster*, was missing in the results from the eDNA analysis; however, this does not necessarily indicate a failure of the eDNA amplification of the newt, as it may be because of an insufficient number of reference sequences, as discussed below.

In physical surveys, the number or composition of species identified in each area is likely to vary among surveys due to factors such as season, weather, and the experience and ability of the surveyors. The results of eDNA surveys should be less variable than in physical surveys because eDNA distribution is less susceptible to these factors. Therefore, eDNA metabarcoding of amphibians may be suitable for large-scale (e.g., national scale) monitoring studies in which the standardization of conditions is necessary.

All five universal primer set candidates tested amplified the 16S rRNA gene fragments from the 16 taxa (with members of all three amphibian orders). Furthermore, Amph16S detected regional intraspecific polymorphisms found in *G. rugosa*, indicating that this primer set with long amplicon length would contribute to revealing intraspecific diversity as well as high taxonomic resolution. In total, eDNA metabarcoding with Amph16S may be used not only for investigating species distribution, but also the genetic diversity of amphibians prone to intraspecific polymorphism (Nishizawa et al. 2011, Tominaga et al. 2013, Oike et al. 2020).

The Japanese fire belly newt, *C*. *pyrrhogaster*, was not detected in the BLAST results. This newt is the only species of the family Salamandridae in this study area and is categorized as “Near Threatened” by the Red List of the Ministry of the Environment. At sampling area J, *C. pyrrhogaster* was detected by the physical surveys, but not by eDNA metabarcoding. However, a sequence with 93% nucleotide similarity and the closest phylogenetic relationship with the *C. pyrrhogaster* 16S rRNA gene (Fig. S4) was found in the eDNA metabarcoding output. A previous study based on a 1407-bp mtDNA sequence (NADH6-tRNAglu-cytb genes) showed that this species contains more than 100 mt haplotypes, and that the intraspecific genetic divergence among major clades of *C. pyrrhogaster* is very large (Tominaga et al. 2013). This large intraspecific variation and lack of 16S rRNA gene data of local *C. pyrrhogaster* populations may have caused the misidentification of this species in the eDNA metabarcoding at site J. In fact, only one 16S rRNA region sequence of this species is available (GenBank: EU880313.1). Our query sequence was undoubtedly a member of Caudata (Fig. S4), and considering the known caudate species in the study site, the sequences is probably of *C. pyrrhogaster* (i.e., Amph16 amplified its DNA). The accumulation of sequences in databases covering local populations is a future challenge and will solve such a problem.

Amphibian populations often have various intra-species lineages and show highly structured spatial-genetic patterns (Nishizawa et al. 2011, Tominaga et al. 2013, Oike et al. 2020). The enhancement of reference sequences is crucial for adequately assigning MOTUs to appropriate taxon (Shaw et al. 2016, Deiner et al. 2017); therefore, collecting representative reference sequences for all populations of each area is desirable and would allow an accurate identification of species of local populations. Although the mitochondrial cytochrome c oxidase I (COI) gene region is generally used for DNA barcoding in animals (Hebert et al. 2003), universal primers for eDNA metabarcoding are often not designed for this region, but for the mitochondrial 16S and 12S rRNA regions (see also Deiner et al. 2017, Miya et al. 2015, Ushio et al. 2017, Valentini et al. 2016). Considering that species-specific detection eDNA assays are mostly designed for protein-coding genes such as cytb, NADH 1, NADH2, and COI (see also Fukumoto et al. 2015, Klymus et al. 2020, Sakata et al. 2017, Schumer et al. 2019), future studies should focus on the whole mitochondrial genome sequences. Although the 16S rRNA region has been recommended as a target for barcoding in amphibians from early stages (Vences et al. 2005), accumulation of reference sequences is still ongoing, especially from the viewpoint of geographic structures, and it must be accelerated to maximize the advantage of high resolution of the new primer herein designed.

While factors such as weather and season can destabilize the results of physical amphibian surveys that involve capture and visual surveillance, eDNA metabarcoding is less susceptible to these factors and can thus provide more stable results. Here, we designed and evaluated some primer sets in the 16S rRNA region, of which there is a relatively rich database of reference sequences available for amphibians. Among them, Amph16S showed relatively good performance in terms of taxonomic resolution and sufficient detectability. eDNA metabarcoding using Amph16S may contribute to rapid surveys of the distribution of amphibian species and possibly to the discovery of new species in the future.

## Supporting information

Supporting Infomation

## Acknowledgements

The State Government of Sarawak permitted us to conduct the project (Permit No. NCCD.907.4.4(JLD.12)-185 and Park Permit No. 436/2015) and to export the collected specimens (Permit No. 16537), and the RDID provided facilities for conducting research. We are grateful to R. ak S. Pungga, P. ak Meleng, and T. Itioka for their support in obtaining permission to conduct research and export specimens. This study was partly supported by JSPS KAKENHI (Grant Numbers: JP16H04735 and JP20H03326), JSPS Core-to-Core Program B to M. Motokawa, Kondo Grant of the Asahi Glass Foundation to K. Nishikawa, JST/JICA, SATREPS to T. Itioka, and Environment Research and Technology Development Fund (Grant Number JPMEERF20164002) to M. Miya.

## Author Contributions

MKS, AK, TN, JK, MM and TM conceived and designed the study. AK, KN and MYH selected and provided appropriate materials. MKS, MUK, AK, TK, RK, MM and TM designed and evaluated the universal primer sets. MN and TS performed the field survey. MKS, MUK, TK, MN, TS and MM performed the laboratory experiments and environmental DNA analysis. MKS and TM wrote and edited the first draft of the manuscript. All authors discussed the results and contributed to the development of the manuscript.

