## Supporting Infomation for "Development and evaluation of PCR primers for environmental DNA (eDNA) metabarcoding of Amphibia"

Supporting Information

Conditions used for downloading sequences from NCBI to create the vertebrate reference database.

The downloads from NCBI were performed according to the following conditions: except for *Homo sapiens* sequences because of their large number: Fish {("Chondrichthyes"[Organism] OR "Dipnoi"[Organism] OR "Actinopterygii"[Organism] OR "Myxini"[Organism] OR "Hyperoartia"[Organism] OR "Coelacanthimorpha"[Organism]) AND (biomol_genomic[PROP] AND ddbj_embl_genbank[filter] AND is_nuccore[filter] AND mitochondrion[filter] AND ("15000"[SLEN] : "30000"[SLEN]))}; Amphibia {"Amphibia"[Organism] AND complete[All Fields] AND (biomol_genomic[PROP] AND ddbj_embl_genbank[filter] AND is_nuccore[filter] AND mitochondrion[filter] AND ("15000"[SLEN] : "30000"[SLEN]))}; Reptile { ("Testudines"[Organism] OR "Lepidosauria"[Organism] OR "Crocodylia"[Organism]) AND (biomol_genomic[PROP] AND ddbj_embl_genbank[filter] AND is_nuccore[filter] AND mitochondrion[filter] AND ("15000"[SLEN] : "30000"[SLEN]))}; Bird { "Aves"[Organism] AND (biomol_genomic[PROP] AND ddbj_embl_genbank[filter] AND is_nuccore[filter] AND mitochondrion[filter] AND ("15000"[SLEN] : "30000"[SLEN]))}; Mammal { "Mammalia"[Organism] NOT ("Homo sapiens"[Organism] OR "Homo sapiens"[All Fields]) AND (biomol_genomic[PROP] AND ddbj_embl_genbank[filter] AND is_nuccore[filter] AND mitochondrion[filter] AND ("15000"[SLEN] : "30000"[SLEN]))}.

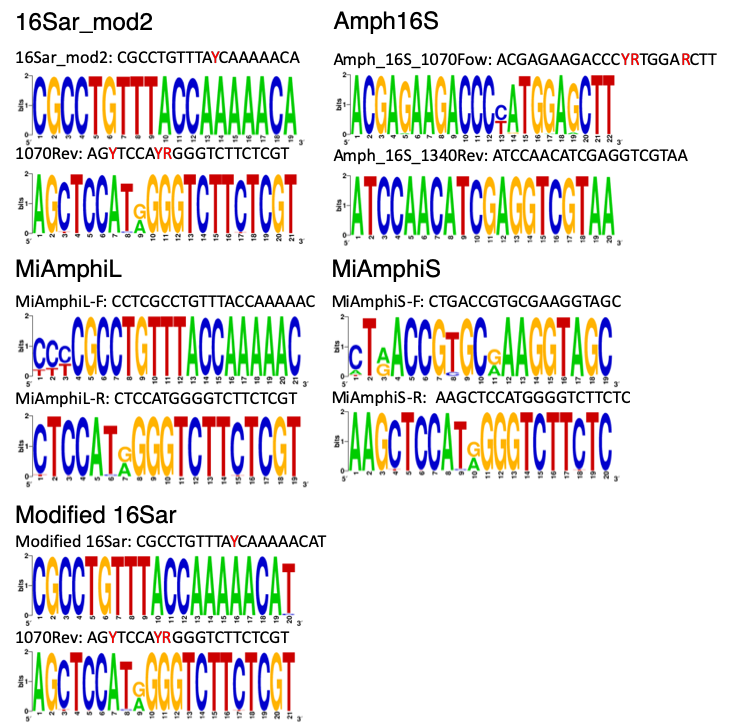

Fig. S1. Sequence logos of the universal primers tested. The size of each base logo is proportional to the percentage of that base in that specific position. The red letter in primer sequence indicates a mixed base.

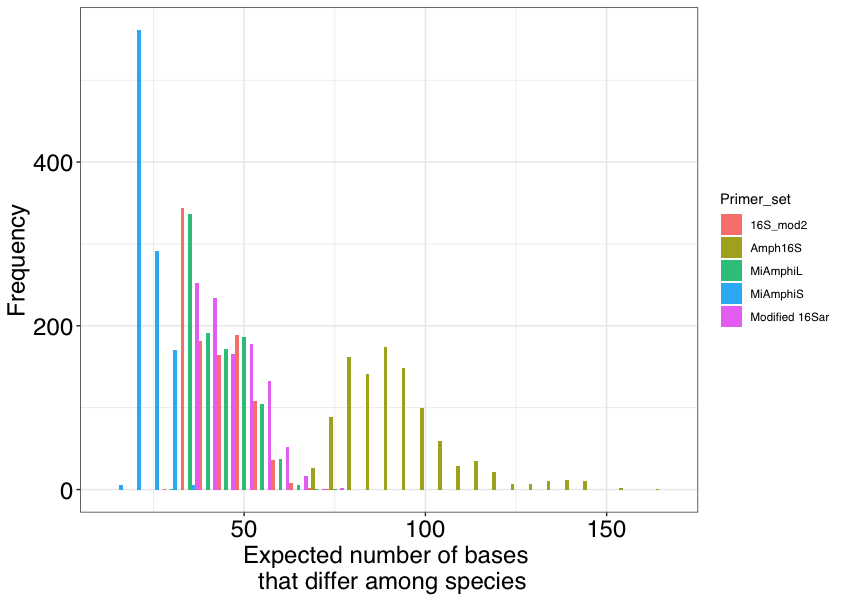

Fig. S2. Distribution of the expected number of bases that differ among species within the amplified region in a primer set. Each category indicates a set of primers; 16Sar_mod2: 16Sar_mod2 and Amph_16S_1070Rev, Amph16S: Amph_16S_1070Fow and Amph_16S_1340Rev, MiAmphiL: MiAmphiL-F and MiAmphiL-R, MiAmphiS: MiAmphiS-F and MiAmphiS-R, Modified 16Sar: Modified 16Sar and Amph_16S_1070Rev.

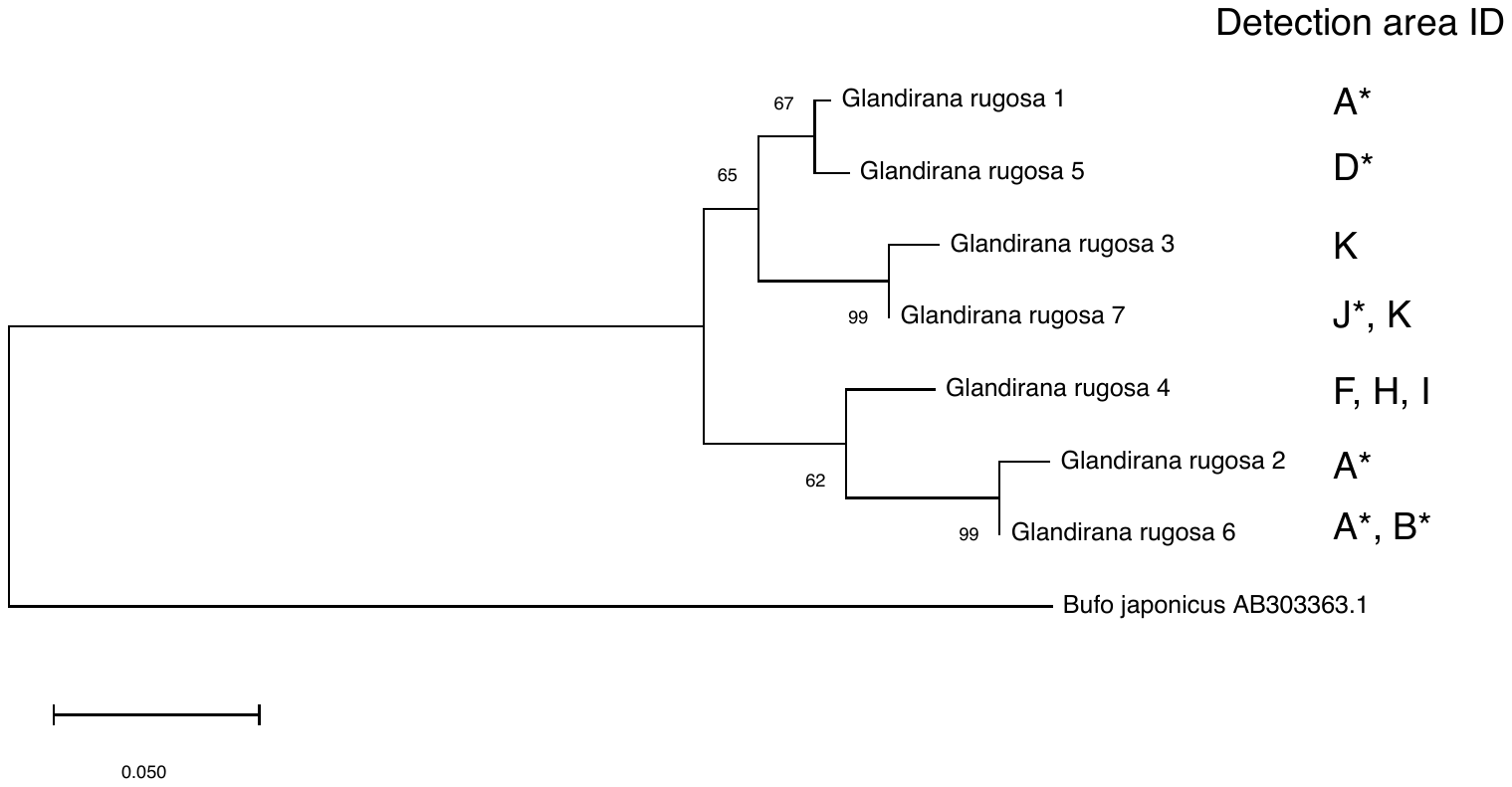

Fig. S3. Maximum likelihood tree of *Glandirana rugosa* on the basis of the 16S rRNA gene using Amph16S (ca. 250bp) with 1,000 bootstrap pseudoreplicates. Bootstrap value is represented at the nodes. Asterisks indicate the areas where *G*. *rugosa* was detected by both eDNA metabarcoding and traditional surveys. The sequence of *Bufo japonicus* is used for the outgroup.

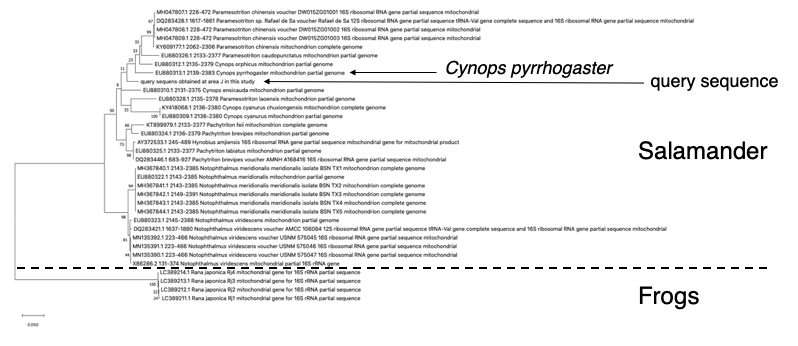

Fig. S4. ML tree of amphibians and an unidentified query sequence at area J. Bootstrap value based on 1,000 pseudoreplicates is represented at the nodes. The query sequence belonged to the salamander group.

Table S1. Amphibian taxa used in primer testing.

| Order | Family | Species | Voucher information |
| --- | --- | --- | --- |
| Anura | Bombinatoridae | *Bombina orientalis* | IABHU Captive individual (no voucher), CBM-DNA 2015-471 |
| Anura | Bufonidae | *Bufo japonicus japonicus* | IABHU Captive individual (no voucher), CBM-DNA 2015-465 |
| Anura | Dicroglossidae | *Fejervarya kawamurai* | CBM-DNA 2015-470 |
| Anura | Hylidae | *Dryophytes japonicus* | IABHU 6123, CBM-DNA 2015-446 |
| Anura | Megophryidae | *Megophrys nasuta* | AKZC SEA01, KUHE 57442 |
| Anura | Microhylidae | *Chaperina fusca* | KUHE 57065 |
| Anura | Microhylidae | *Kalophrynus meizon* | KUHE 57154 |
| Anura | Microhylidae | *Microhyla malang* | KUHE 57066 |
| Anura | Pipidae | *Xenopus laevis* | CBM-DNA 2015-472 |
| Anura | Ranidae | *Pelophylax nigromaculatus* | CBM-DNA 2015-468 |
| Anura | Rhacophoridae | *Buergeria buergeri* | CBM-DNA 2015-467 |
| Anura | Rhacophoridae | *Rhacophorus nigropalmatus* | KUHE 57173 |
| Anura | Scaphiopodidae | *Scaphiopus holbrookii* | CBM-DNA 2015-475 |
| Gymnophiona | Ichthyophiidae | *Ichthyophis biangularis* | KUHE 57201 |
| Caudata | Salamandridae | *Cynops pyrrhogaster* | IABHU Captive individual (no voucher), CBM-DNA 2015-473 |
| Caudata | Hynobiidae | *Hynobius naevius* | IABHU Captive individual (no voucher), CBM-DNA 2015-474 |

IABHU: Institute for Amphibian Biology, Hiroshima University (now called Amphibian Research Center, Hiroshima University).

AKZC: AK's Zoological Collection. AK: AK's field number of tissue specimens.

KUHE: Graduate School of Human and Environmental Studies, Kyoto University

CBM-DNA: Tissue collection, Natural History Museum and Institute Chiba (without voucher specimens)

Table S2. The list of each site information.

| Site ID | Latitude (N) | Longitude (E) | Physical Survey |
| --- | --- | --- | --- |
| A01 | 36.66 | 139.93 | + |
| A02 | 36.65 | 139.93 | + |
| A03 | 36.65 | 139.93 | + |
| A04 | 36.65 | 139.93 | + |
| A05 | 36.65 | 139.93 | + |
| A06 | 36.66 | 139.93 | + |
| A07 | 36.65 | 139.93 | + |
| A08 | 36.66 | 139.93 | + |
| A09 | 36.65 | 139.93 | + |
| A10 | 36.64 | 139.93 | + |
| A11 | 36.66 | 139.93 | - |
| A12 | 36.67 | 139.94 | - |
| A13 | 36.64 | 139.95 | - |
| A14 | 36.66 | 139.93 | - |
| B01 | 39.10 | 141.00 | - |
| B02 | 39.09 | 141.02 | + |
| B03 | 39.09 | 141.03 | + |
| B04 | 39.09 | 141.04 | - |
| B05 | 39.09 | 141.04 | - |
| B06 | 39.08 | 141.05 | + |
| B07 | 39.08 | 141.05 | + |
| B08 | 39.08 | 141.04 | - |
| B09 | 39.08 | 141.04 | - |
| B10 | 39.08 | 141.04 | + |
| B11 | 39.08 | 141.05 | + |
| C01 | 35.90 | 140.49 | - |
| C02 | 35.83 | 140.49 | - |
| C03 | 35.83 | 140.49 | - |
| C04 | 35.72 | 140.48 | - |
| C05 | 35.70 | 140.47 | + |
| C06 | 35.71 | 140.49 | + |
| C07 | 35.71 | 140.48 | + |
| C08 | 35.71 | 140.48 | + |
| C09 | 35.72 | 140.49 | + |
| C10 | 35.72 | 140.48 | + |
| C11 | 35.71 | 140.48 | - |
| C12 | 35.70 | 140.48 | + |
| C13 | 35.70 | 140.48 | + |
| C14 | 35.70 | 140.47 | + |
| C15 | 35.70 | 140.48 | + |
| C16 | 35.70 | 140.48 | + |
| C17 | 35.70 | 140.48 | + |
| C18 | 35.71 | 140.48 | + |
| C19 | 35.70 | 140.47 | + |
| C20 | 35.70 | 140.47 | - |
| D01 | 36.74 | 136.96 | - |
| D02 | 36.68 | 136.93 | - |
| D03 | 36.68 | 136.93 | + |
| D04 | 36.68 | 136.93 | + |
| D05 | 36.67 | 136.93 | + |
| D06 | 36.67 | 136.93 | + |
| D07 | 36.66 | 136.93 | + |
| D08 | 36.66 | 136.93 | + |
| D09 | 36.66 | 136.93 | - |
| D10 | 36.65 | 136.94 | - |
| D11 | 36.68 | 136.90 | - |
| D12 | 36.67 | 136.90 | - |
| D13 | 36.67 | 136.90 | + |
| D14 | 36.67 | 136.91 | + |
| D15 | 36.66 | 136.91 | + |
| D16 | 36.65 | 136.92 | + |
| D17 | 36.64 | 136.92 | + |
| D18 | 36.64 | 136.93 | + |
| D19 | 36.64 | 136.93 | - |
| D20 | 36.59 | 136.99 | - |
| F02 | 34.57 | 136.56 | + |
| F03 | 34.57 | 136.56 | + |
| F04 | 34.57 | 136.56 | + |
| F05 | 34.57 | 136.56 | + |
| F10 | 34.57 | 136.56 | + |
| F11 | 34.57 | 136.56 | + |
| F12 | 34.57 | 136.56 | + |
| F18 | 34.56 | 136.57 | + |
| F27 | 34.56 | 136.56 | + |
| F29 | 34.56 | 136.57 | + |
| F33 | 34.57 | 136.56 | - |
| F34 | 34.53 | 136.57 | - |
| G01 | 35.09 | 136.62 | + |
| G02 | 35.09 | 136.62 | + |
| G03 | 35.09 | 136.62 | + |
| G04 | 35.09 | 136.62 | + |
| G06 | 35.09 | 136.62 | + |
| G07 | 35.09 | 136.62 | + |
| G09 | 35.09 | 136.62 | + |
| G10 | 35.09 | 136.62 | + |
| G12 | 35.09 | 136.62 | + |
| G14 | 35.09 | 136.62 | + |
| G15 | 35.09 | 136.62 | + |
| G16 | 35.09 | 136.62 | + |
| G17 | 35.09 | 136.62 | + |
| G19 | 35.09 | 136.62 | + |
| G21 | 35.09 | 136.62 | + |
| G22 | 35.09 | 136.62 | + |
| G25 | 35.09 | 136.62 | + |
| G26 | 35.08 | 136.63 | + |
| G28 | 35.08 | 136.63 | + |
| G29 | 35.09 | 136.62 | - |
| H01 | 35.51 | 136.22 | + |
| H02 | 35.51 | 136.21 | + |
| H03 | 35.51 | 136.21 | + |
| H04 | 35.51 | 136.21 | + |
| H05 | 35.51 | 136.22 | + |
| H06 | 35.51 | 136.22 | + |
| H07 | 35.52 | 136.21 | - |
| H08 | 35.51 | 136.21 | - |
| H09 | 35.51 | 136.21 | - |
| H10 | 35.51 | 136.22 | - |
| I01 | 34.91 | 135.77 | + |
| I02 | 34.91 | 135.77 | + |
| I03 | 34.90 | 135.77 | + |
| I04 | 34.90 | 135.76 | + |
| I05 | 34.91 | 135.77 | + |
| I06 | 34.91 | 135.77 | + |
| I07 | 34.91 | 135.76 | + |
| I08 | 34.90 | 135.75 | + |
| I09 | 34.91 | 135.79 | - |
| I10 | 34.91 | 135.77 | - |
| I11 | 34.90 | 135.74 | - |
| I12 | 34.91 | 135.78 | - |
| J01 | 34.46 | 131.68 | + |
| J02 | 34.47 | 131.67 | - |
| J03 | 34.47 | 131.67 | + |
| J04 | 34.47 | 131.67 | - |
| J05 | 34.47 | 131.68 | - |
| J06 | 34.47 | 131.69 | - |
| J09 | 34.47 | 131.66 | + |
| J10 | 34.47 | 131.66 | + |
| J11 | 34.47 | 131.66 | + |
| J12 | 34.47 | 131.65 | + |
| J13 | 34.47 | 131.66 | + |
| J14 | 34.47 | 131.66 | + |
| J19 | 34.47 | 131.67 | + |
| J20 | 34.47 | 131.66 | + |
| J21 | 34.47 | 131.66 | + |
| J22 | 34.46 | 131.66 | + |
| J23 | 34.47 | 131.66 | + |
| J24 | 34.47 | 131.66 | + |
| J25 | 34.47 | 131.67 | + |
| J32 | 34.47 | 131.66 | + |
| K01 | 33.28 | 130.28 | + |
| K02 | 33.28 | 130.27 | + |
| K03 | 33.28 | 130.28 | + |
| K04 | 33.28 | 130.27 | + |
| K05 | 33.28 | 130.27 | + |
| K06 | 33.28 | 130.28 | + |
| K07 | 33.28 | 130.28 | + |
| K08 | 33.29 | 130.28 | + |
| K09 | 33.29 | 130.28 | + |
| K10 | 33.29 | 130.28 | + |
| K11 | 33.28 | 130.27 | + |
| K12 | 33.28 | 130.26 | + |
| K13 | 33.28 | 130.26 | + |
| K14 | 33.28 | 130.26 | + |
| K15 | 33.28 | 130.26 | + |
| K16 | 33.27 | 130.26 | + |
| K17 | 33.28 | 130.28 | + |
| K18 | 33.29 | 130.28 | + |
| K19 | 33.29 | 130.26 | + |
| K20 | 33.29 | 130.28 | + |
| K21 | 33.29 | 130.26 | + |

In column “Physical surveys”, “+” indicates that it was performed, and “-” indicates not performed.

Table S3. The results of in silico PCR.

| maxdiff = 0 |  |  |  |  |  |
| --- | --- | --- | --- | --- | --- |
| Primer set / Reference taxa | Fish | Amphibia | Reptile | Bird | Mammal |
| MiAmphiL | 0.2% | 3.1% | 0% | 0% | 0% |
| MiAmphiS | 0% | 5.0% | 0% | 0% | 0.0% |
| Modified 16Sar | 62.6% | 89.9% | 0.7% | 0% | 96.4% |
| 16Sar_mod2 | 66.9% | 94.9% | 3.9% | 0% | 96.6% |
| Amph16S | 64.9% | 96.4% | 56.1% | 97.6% | 96.4% |
| maxdiff = 1 |  |  |  |  |  |
| Primer set / Reference taxa | Fish | Amphibia | Reptile | Bird | Mammal |
| MiAmphiL | 48.4% | 75.5% | 0.1% | 0% | 89.4% |
| MiAmphiS | 32.9% | 52.5% | 0% | 0.0% | 75.8% |
| Modified 16Sar | 99.0% | 97.6% | 38.3% | 83.8% | 98.1% |
| 16Sar_mod2 | 99.1% | 97.6% | 46.8% | 84.4% | 98.1% |
| Amph16S | 97.0% | 98.6% | 69.1% | 99.3% | 98.0% |
| maxdiff = 2 |  |  |  |  |  |
| Primer set / Reference taxa | Fish | Amphibia | Reptile | Bird | Mammal |
| MiAmphiL | 98.3% | 93.6% | 8.2% | 0% | 96.9% |
| MiAmphiS | 91.5% | 96.2% | 28.9% | 4.9% | 96.8% |
| Modified 16Sar | 99.3% | 97.6% | 73.9% | 99.1% | 98.2% |
| 16Sar_mod2 | 99.4% | 97.7% | 74.3% | 99.1% | 98.3% |
| Amph16S | 97.2% | 98.7% | 90.3% | 99.4% | 98.1% |

The “maxdiff” indicates that maximum number of mismatches allowed between each primer sequence and the reference sequence. The percentage indicates the number of sequences determined to be amplified potentially among the reference sequences of each taxon (Fish: 7861, Amphibia: 959, Reptile: 1170, Bird: 2988, Mammal: 13118).

Table S4. Result of eDNA metabarcoding.

| Site ID | *Buergeria buergeri* | *Bufo japonicus* | *Bu. torrenticola* | *Fejervarya limnocharis* | *Glandirana rugosa* | *Dryophytes japonica* | *Lithobates catesbeianus* | *Pelophylax* sp. | *P. nigromaculatus* | *P. porosus* subsp. | *Rana japonica* | *Ra. ornativentris* | *Ra. tagoi tagoi* | *Zhangixalus*  *arboreus* | *Z.. schlegelii* |
| --- | --- | --- | --- | --- | --- | --- | --- | --- | --- | --- | --- | --- | --- | --- | --- |
| A01 | 0 | 0 | 0 | 0 | 118 | 4669 | 0 | 0 | 0 | 4430 | 89 | 0 | 0 | 0 | 0 |
| A02 | 0 | 0 | 0 | 0 | 0 | 6623 | 0 | 0 | 0 | 38781 | 0 | 0 | 0 | 0 | 0 |
| A03 | 0 | 0 | 0 | 0 | 1341 | 4377 | 0 | 0 | 0 | 6866 | 175 | 0 | 0 | 0 | 0 |
| A04 | 0 | 0 | 0 | 0 | 591 | 6764 | 0 | 0 | 0 | 41624 | 10 | 0 | 0 | 0 | 0 |
| A05 | 0 | 0 | 0 | 0 | 5409 | 6163 | 0 | 0 | 0 | 15361 | 174 | 0 | 0 | 0 | 0 |
| A06 | 0 | 86 | 0 | 0 | 711 | 6373 | 0 | 0 | 0 | 1766 | 90 | 0 | 0 | 0 | 0 |
| A07 | 0 | 0 | 0 | 0 | 248 | 6921 | 0 | 0 | 0 | 36799 | 134 | 0 | 0 | 0 | 0 |
| A08 | 0 | 0 | 0 | 0 | 1163 | 2292 | 0 | 0 | 0 | 7798 | 260 | 0 | 0 | 0 | 0 |
| A09 | 0 | 0 | 0 | 0 | 1256 | 3870 | 0 | 0 | 0 | 14118 | 47 | 0 | 0 | 0 | 0 |
| A10 | 0 | 0 | 0 | 0 | 7186 | 2123 | 0 | 0 | 0 | 8470 | 219 | 0 | 0 | 0 | 0 |
| A11 | 0 | 0 | 0 | 0 | 2605 | 1988 | 0 | 0 | 0 | 3627 | 247 | 0 | 0 | 0 | 0 |
| A12 | 0 | 0 | 0 | 0 | 1444 | 1185 | 0 | 0 | 0 | 3541 | 326 | 0 | 0 | 0 | 0 |
| A13 | 0 | 0 | 0 | 0 | 1150 | 1618 | 0 | 0 | 0 | 3415 | 315 | 0 | 0 | 0 | 0 |
| A14 | 0 | 0 | 0 | 0 | 0 | 12469 | 0 | 0 | 0 | 0 | 0 | 0 | 0 | 0 | 0 |
| B01 | 0 | 0 | 0 | 0 | 95 | 229 | 0 | 0 | 0 | 1304 | 0 | 0 | 0 | 0 | 59 |
| B02 | 0 | 0 | 0 | 0 | 135 | 700 | 0 | 0 | 0 | 2606 | 0 | 0 | 0 | 0 | 30 |
| B03 | 0 | 0 | 0 | 0 | 0 | 467 | 0 | 0 | 0 | 547 | 0 | 0 | 0 | 0 | 40 |
| B04 | 0 | 0 | 0 | 0 | 322 | 753 | 0 | 0 | 0 | 2611 | 0 | 0 | 0 | 0 | 52 |
| B05 | 0 | 0 | 0 | 0 | 151 | 415 | 0 | 0 | 0 | 2162 | 0 | 0 | 0 | 0 | 28 |
| B06 | 0 | 0 | 0 | 0 | 301 | 1399 | 0 | 0 | 0 | 3932 | 0 | 0 | 117 | 0 | 0 |
| B07 | 0 | 0 | 0 | 0 | 1448 | 14694 | 0 | 0 | 0 | 53264 | 0 | 0 | 0 | 0 | 1001 |
| B08 | 0 | 0 | 0 | 0 | 858 | 621 | 0 | 0 | 0 | 1164 | 0 | 0 | 0 | 0 | 0 |
| B09 | 0 | 0 | 0 | 0 | 116 | 805 | 0 | 0 | 0 | 834 | 0 | 30 | 0 | 0 | 117 |
| B10 | 0 | 0 | 0 | 0 | 949 | 871 | 0 | 0 | 0 | 3001 | 0 | 0 | 0 | 0 | 0 |
| B11 | 0 | 0 | 0 | 0 | 333 | 1118 | 0 | 0 | 0 | 7477 | 0 | 0 | 0 | 0 | 33 |
| C01 | 0 | 0 | 0 | 0 | 0 | 0 | 0 | 0 | 0 | 0 | 0 | 0 | 0 | 0 | 0 |
| C02 | 0 | 0 | 0 | 0 | 0 | 2750 | 0 | 0 | 0 | 0 | 176 | 0 | 0 | 0 | 57 |
| C03 | 0 | 0 | 0 | 0 | 0 | 280 | 0 | 0 | 0 | 0 | 0 | 0 | 0 | 0 | 0 |
| C04 | 0 | 0 | 0 | 197 | 0 | 1560 | 36 | 0 | 0 | 658 | 540 | 0 | 0 | 0 | 21 |
| C05 | 0 | 0 | 0 | 0 | 0 | 0 | 0 | 0 | 0 | 0 | 0 | 0 | 0 | 0 | 0 |
| C06 | 0 | 0 | 0 | 0 | 0 | 4968 | 80 | 0 | 0 | 1448 | 0 | 0 | 0 | 0 | 22 |
| C07 | 0 | 0 | 0 | 0 | 0 | 4027 | 0 | 0 | 0 | 917 | 0 | 0 | 0 | 0 | 0 |
| C08 | 0 | 0 | 0 | 0 | 0 | 546 | 0 | 0 | 0 | 536 | 0 | 0 | 0 | 0 | 0 |
| C09 | 0 | 0 | 0 | 0 | 0 | 24 | 0 | 0 | 0 | 33 | 11 | 0 | 0 | 0 | 0 |
| C10 | 0 | 0 | 0 | 0 | 0 | 1923 | 0 | 0 | 0 | 159 | 0 | 0 | 0 | 0 | 0 |
| C11 | 0 | 0 | 0 | 0 | 0 | 543 | 0 | 0 | 0 | 256 | 356 | 0 | 0 | 0 | 26 |
| C12 | 0 | 0 | 0 | 0 | 0 | 5136 | 0 | 0 | 0 | 6054 | 85 | 0 | 0 | 0 | 0 |
| C13 | 0 | 0 | 0 | 0 | 0 | 1871 | 0 | 0 | 0 | 16814 | 14 | 0 | 0 | 0 | 0 |
| C14 | 0 | 0 | 0 | 0 | 0 | 638 | 0 | 0 | 0 | 25955 | 0 | 0 | 0 | 0 | 0 |
| C15 | 0 | 0 | 0 | 0 | 0 | 8638 | 0 | 0 | 0 | 0 | 35 | 0 | 0 | 0 | 0 |
| C16 | 0 | 0 | 0 | 0 | 0 | 0 | 0 | 0 | 0 | 347 | 0 | 0 | 0 | 0 | 0 |
| C17 | 0 | 0 | 0 | 0 | 0 | 3563 | 0 | 0 | 0 | 1214 | 19 | 0 | 0 | 0 | 22 |
| C18 | 0 | 0 | 0 | 0 | 0 | 3265 | 0 | 0 | 0 | 4515 | 0 | 0 | 0 | 0 | 0 |
| C19 | 0 | 0 | 0 | 0 | 0 | 2207 | 344 | 0 | 0 | 3556 | 0 | 0 | 0 | 0 | 0 |
| C20 | 0 | 0 | 0 | 0 | 0 | 1516 | 182 | 0 | 0 | 776 | 284 | 0 | 0 | 0 | 0 |
| D01 | 0 | 0 | 0 | 0 | 48 | 146 | 0 | 0 | 0 | 0 | 0 | 0 | 0 | 0 | 0 |
| D02 | 0 | 0 | 0 | 0 | 2842 | 35 | 0 | 0 | 620 | 0 | 0 | 0 | 0 | 0 | 0 |
| D03 | 0 | 0 | 0 | 0 | 694 | 288 | 0 | 0 | 2206 | 0 | 0 | 0 | 0 | 0 | 0 |
| D04 | 0 | 0 | 0 | 0 | 2161 | 178 | 0 | 0 | 5177 | 0 | 0 | 0 | 0 | 0 | 0 |
| D05 | 0 | 0 | 0 | 0 | 1263 | 95 | 0 | 0 | 35902 | 0 | 0 | 0 | 0 | 0 | 0 |
| D06 | 0 | 0 | 0 | 0 | 1177 | 0 | 0 | 0 | 7066 | 0 | 0 | 0 | 0 | 0 | 0 |
| D07 | 0 | 0 | 119 | 0 | 1026 | 383 | 0 | 0 | 18669 | 0 | 0 | 0 | 0 | 0 | 0 |
| D08 | 0 | 0 | 0 | 0 | 1910 | 372 | 0 | 0 | 11639 | 0 | 0 | 0 | 0 | 0 | 0 |
| D09 | 0 | 0 | 0 | 0 | 824 | 0 | 0 | 0 | 845 | 0 | 0 | 0 | 0 | 0 | 0 |
| D10 | 0 | 0 | 0 | 0 | 0 | 0 | 0 | 0 | 0 | 952 | 0 | 0 | 0 | 0 | 0 |
| D11 | 0 | 0 | 0 | 0 | 1321 | 0 | 0 | 0 | 978 | 0 | 0 | 0 | 0 | 0 | 0 |
| D12 | 0 | 0 | 0 | 0 | 1759 | 0 | 0 | 0 | 1863 | 0 | 0 | 0 | 0 | 0 | 0 |
| D13 | 0 | 0 | 0 | 0 | 1515 | 0 | 0 | 0 | 828 | 0 | 0 | 0 | 0 | 0 | 0 |
| D14 | 0 | 0 | 0 | 0 | 1752 | 0 | 0 | 0 | 5859 | 0 | 0 | 0 | 0 | 0 | 0 |
| D15 | 0 | 0 | 0 | 0 | 2189 | 0 | 0 | 0 | 2507 | 0 | 0 | 0 | 0 | 0 | 0 |
| D16 | 0 | 0 | 0 | 0 | 2692 | 0 | 0 | 0 | 7241 | 0 | 0 | 0 | 0 | 0 | 0 |
| D17 | 0 | 0 | 0 | 0 | 373 | 0 | 0 | 0 | 0 | 0 | 0 | 0 | 0 | 0 | 0 |
| D18 | 0 | 0 | 0 | 0 | 1631 | 2954 | 0 | 0 | 13477 | 0 | 0 | 0 | 0 | 0 | 0 |
| D19 | 0 | 0 | 0 | 0 | 0 | 913 | 0 | 0 | 4205 | 0 | 0 | 0 | 0 | 0 | 0 |
| D20 | 0 | 0 | 0 | 0 | 0 | 0 | 0 | 0 | 0 | 0 | 0 | 0 | 0 | 0 | 0 |
| F02 | 0 | 0 | 0 | 3294 | 0 | 251 | 0 | 0 | 25734 | 0 | 0 | 0 | 0 | 0 | 0 |
| F03 | 0 | 0 | 0 | 7687 | 0 | 775 | 244 | 0 | 32696 | 0 | 0 | 0 | 0 | 0 | 0 |
| F04 | 0 | 0 | 0 | 8230 | 0 | 237 | 150 | 0 | 35628 | 0 | 0 | 0 | 0 | 0 | 0 |
| F05 | 0 | 0 | 0 | 37709 | 0 | 569 | 0 | 0 | 4149 | 0 | 0 | 0 | 0 | 0 | 0 |
| F10 | 0 | 0 | 0 | 181 | 0 | 470 | 140 | 0 | 14427 | 0 | 0 | 0 | 0 | 0 | 0 |
| F11 | 0 | 0 | 0 | 3285 | 0 | 3693 | 0 | 0 | 40307 | 0 | 0 | 0 | 0 | 0 | 0 |
| F12 | 0 | 0 | 0 | 61551 | 0 | 126 | 16 | 0 | 2374 | 75 | 0 | 0 | 0 | 0 | 0 |
| F18 | 0 | 0 | 0 | 9413 | 595 | 2219 | 1386 | 0 | 18901 | 879 | 0 | 0 | 0 | 0 | 0 |
| F27 | 0 | 0 | 0 | 12559 | 0 | 647 | 306 | 0 | 9100 | 0 | 0 | 0 | 0 | 0 | 0 |
| F29 | 0 | 0 | 0 | 2874 | 0 | 1131 | 0 | 0 | 16193 | 0 | 0 | 0 | 0 | 0 | 0 |
| F33 | 0 | 0 | 0 | 4343 | 0 | 406 | 8354 | 0 | 17665 | 0 | 0 | 0 | 0 | 0 | 0 |
| F34 | 0 | 0 | 0 | 263 | 897 | 87 | 109 | 0 | 2264 | 0 | 0 | 0 | 0 | 0 | 0 |
| G01 | 0 | 0 | 0 | 2207 | 0 | 13 | 4253 | 0 | 9492 | 0 | 0 | 0 | 0 | 33 | 0 |
| G02 | 0 | 0 | 0 | 875 | 0 | 54 | 3695 | 0 | 14553 | 0 | 0 | 0 | 0 | 27 | 0 |
| G03 | 0 | 0 | 0 | 2384 | 0 | 0 | 3456 | 0 | 14254 | 0 | 0 | 0 | 0 | 23 | 0 |
| G04 | 0 | 0 | 0 | 3799 | 0 | 0 | 1256 | 0 | 18348 | 0 | 0 | 0 | 0 | 0 | 0 |
| G06 | 0 | 0 | 0 | 13430 | 0 | 0 | 1498 | 0 | 4681 | 0 | 0 | 0 | 0 | 13 | 0 |
| G07 | 0 | 0 | 0 | 1214 | 0 | 85 | 0 | 0 | 41397 | 0 | 0 | 0 | 0 | 0 | 0 |
| G09 | 0 | 0 | 0 | 2006 | 0 | 210 | 0 | 0 | 41200 | 0 | 0 | 0 | 0 | 0 | 0 |
| G10 | 0 | 0 | 0 | 0 | 0 | 0 | 0 | 0 | 47009 | 0 | 0 | 0 | 0 | 0 | 0 |
| G12 | 0 | 0 | 0 | 232 | 0 | 0 | 1327 | 0 | 6259 | 0 | 0 | 0 | 0 | 0 | 0 |
| G14 | 0 | 0 | 0 | 0 | 0 | 0 | 4708 | 0 | 460 | 0 | 0 | 0 | 0 | 0 | 0 |
| G15 | 0 | 0 | 0 | 0 | 0 | 0 | 7464 | 0 | 72 | 0 | 0 | 0 | 0 | 0 | 0 |
| G16 | 0 | 0 | 0 | 0 | 0 | 0 | 3992 | 0 | 537 | 0 | 0 | 0 | 0 | 0 | 0 |
| G17 | 0 | 0 | 0 | 0 | 0 | 0 | 1486 | 0 | 155 | 0 | 0 | 0 | 0 | 92 | 0 |
| G19 | 0 | 0 | 0 | 3822 | 0 | 0 | 0 | 0 | 2972 | 0 | 0 | 0 | 0 | 0 | 0 |
| G21 | 0 | 0 | 0 | 849 | 0 | 0 | 0 | 0 | 0 | 0 | 0 | 0 | 0 | 200 | 0 |
| G22 | 0 | 0 | 0 | 0 | 0 | 0 | 4487 | 0 | 29051 | 0 | 0 | 0 | 0 | 3390 | 0 |
| G25 | 0 | 0 | 0 | 0 | 0 | 0 | 6905 | 0 | 35825 | 0 | 0 | 0 | 0 | 4723 | 0 |
| G26 | 0 | 0 | 0 | 2031 | 0 | 2537 | 0 | 0 | 28119 | 0 | 0 | 0 | 0 | 0 | 0 |
| G28 | 0 | 0 | 0 | 9553 | 0 | 918 | 0 | 0 | 44087 | 0 | 0 | 0 | 0 | 0 | 0 |
| G29 | 0 | 0 | 0 | 1575 | 0 | 14 | 3219 | 0 | 10290 | 0 | 0 | 0 | 0 | 0 | 0 |
| H01 | 0 | 0 | 0 | 0 | 900 | 0 | 369 | 0 | 7635 | 9098 | 0 | 0 | 0 | 0 | 0 |
| H02 | 0 | 0 | 0 | 0 | 74 | 0 | 164 | 0 | 3367 | 165 | 0 | 0 | 0 | 0 | 0 |
| H03 | 0 | 0 | 0 | 0 | 825 | 192 | 0 | 0 | 779 | 1595 | 0 | 0 | 213 | 0 | 0 |
| H04 | 0 | 0 | 0 | 0 | 0 | 0 | 0 | 0 | 1309 | 0 | 0 | 0 | 0 | 0 | 0 |
| H05 | 0 | 0 | 0 | 0 | 0 | 0 | 0 | 0 | 2206 | 432 | 0 | 0 | 32 | 32 | 0 |
| H06 | 0 | 0 | 0 | 0 | 0 | 0 | 0 | 0 | 909 | 38 | 0 | 151 | 0 | 0 | 0 |
| H07 | 0 | 0 | 0 | 0 | 165 | 0 | 37 | 0 | 2500 | 856 | 0 | 0 | 0 | 0 | 0 |
| H08 | 0 | 0 | 0 | 0 | 0 | 0 | 0 | 0 | 1281 | 184 | 0 | 0 | 0 | 0 | 17 |
| H09 | 0 | 0 | 0 | 0 | 0 | 0 | 0 | 0 | 1139 | 0 | 0 | 0 | 0 | 0 | 0 |
| H10 | 0 | 0 | 0 | 0 | 161 | 0 | 79 | 0 | 1414 | 18324 | 0 | 0 | 117 | 0 | 0 |
| I01 | 0 | 0 | 0 | 2771 | 106 | 0 | 0 | 0 | 0 | 0 | 0 | 0 | 0 | 0 | 0 |
| I02 | 0 | 0 | 0 | 869 | 562 | 0 | 0 | 0 | 0 | 0 | 0 | 0 | 0 | 0 | 0 |
| I03 | 0 | 0 | 0 | 1067 | 466 | 0 | 0 | 0 | 0 | 0 | 0 | 0 | 0 | 0 | 0 |
| I04 | 0 | 0 | 0 | 576 | 115 | 0 | 0 | 0 | 0 | 0 | 0 | 0 | 0 | 0 | 0 |
| I05 | 0 | 0 | 0 | 734 | 588 | 0 | 0 | 0 | 307 | 0 | 0 | 0 | 0 | 0 | 0 |
| I06 | 0 | 0 | 0 | 4823 | 1924 | 65 | 0 | 0 | 4318 | 0 | 0 | 0 | 0 | 0 | 0 |
| I07 | 0 | 0 | 0 | 7659 | 8536 | 0 | 0 | 0 | 0 | 0 | 0 | 0 | 0 | 0 | 0 |
| I08 | 0 | 0 | 0 | 15488 | 1845 | 0 | 0 | 0 | 0 | 0 | 0 | 0 | 0 | 0 | 0 |
| I09 | 0 | 0 | 0 | 0 | 0 | 0 | 0 | 0 | 0 | 0 | 0 | 0 | 0 | 0 | 0 |
| I10 | 0 | 0 | 0 | 0 | 0 | 0 | 0 | 0 | 0 | 0 | 0 | 0 | 0 | 0 | 0 |
| I11 | 0 | 0 | 0 | 9001 | 0 | 0 | 0 | 0 | 0 | 0 | 0 | 0 | 0 | 0 | 0 |
| I12 | 0 | 0 | 0 | 10 | 43 | 0 | 0 | 0 | 0 | 0 | 0 | 0 | 0 | 0 | 0 |
| J01 | 0 | 0 | 0 | 0 | 1464 | 125 | 0 | 0 | 779 | 0 | 435 | 0 | 0 | 0 | 0 |
| J02 | 0 | 0 | 0 | 0 | 10066 | 115 | 0 | 0 | 1583 | 0 | 4235 | 0 | 0 | 0 | 0 |
| J03 | 0 | 0 | 0 | 0 | 1100 | 49 | 0 | 0 | 87 | 0 | 271 | 0 | 0 | 0 | 0 |
| J04 | 0 | 0 | 0 | 0 | 419 | 29 | 0 | 0 | 0 | 0 | 0 | 0 | 0 | 0 | 0 |
| J05 | 0 | 0 | 0 | 0 | 8048 | 0 | 0 | 0 | 0 | 0 | 495 | 0 | 0 | 92 | 169 |
| J06 | 0 | 0 | 0 | 0 | 2764 | 0 | 0 | 0 | 0 | 0 | 0 | 0 | 0 | 0 | 0 |
| J09 | 0 | 0 | 0 | 0 | 152 | 670 | 0 | 0 | 153 | 0 | 0 | 0 | 0 | 0 | 0 |
| J10 | 0 | 0 | 0 | 0 | 0 | 0 | 0 | 0 | 0 | 0 | 0 | 0 | 0 | 0 | 0 |
| J11 | 0 | 0 | 0 | 0 | 550 | 0 | 0 | 0 | 0 | 0 | 0 | 0 | 248 | 0 | 0 |
| J12 | 0 | 0 | 0 | 0 | 373 | 0 | 0 | 0 | 0 | 0 | 0 | 0 | 0 | 0 | 0 |
| J13 | 0 | 0 | 0 | 0 | 640 | 73 | 0 | 0 | 0 | 0 | 94 | 0 | 0 | 0 | 0 |
| J14 | 0 | 0 | 0 | 0 | 0 | 154 | 0 | 0 | 832 | 0 | 0 | 0 | 0 | 0 | 0 |
| J19 | 0 | 0 | 0 | 0 | 2840 | 484 | 0 | 0 | 5223 | 0 | 1912 | 0 | 152 | 0 | 22 |
| J20 | 0 | 0 | 0 | 0 | 1970 | 0 | 0 | 0 | 9007 | 0 | 0 | 0 | 0 | 0 | 0 |
| J21 | 0 | 0 | 0 | 0 | 3065 | 0 | 0 | 0 | 1327 | 0 | 54 | 0 | 0 | 0 | 0 |
| J22 | 0 | 0 | 0 | 0 | 3782 | 0 | 0 | 0 | 0 | 0 | 303 | 0 | 0 | 0 | 0 |
| J23 | 0 | 0 | 0 | 0 | 3509 | 0 | 0 | 0 | 5762 | 0 | 0 | 0 | 0 | 0 | 0 |
| J24 | 0 | 0 | 0 | 0 | 664 | 0 | 0 | 0 | 0 | 0 | 117 | 0 | 0 | 0 | 0 |
| J25 | 0 | 0 | 0 | 0 | 2216 | 190 | 0 | 0 | 9420 | 0 | 1683 | 0 | 0 | 0 | 0 |
| J32 | 0 | 0 | 0 | 0 | 1150 | 203 | 0 | 0 | 1197 | 0 | 15 | 0 | 0 | 0 | 0 |
| K01 | 0 | 0 | 0 | 3021 | 0 | 30 | 2048 | 0 | 0 | 0 | 36 | 0 | 0 | 0 | 0 |
| K02 | 0 | 0 | 0 | 3970 | 96 | 0 | 57 | 79 | 0 | 0 | 0 | 0 | 0 | 0 | 0 |
| K03 | 0 | 0 | 0 | 6089 | 0 | 0 | 511 | 72 | 0 | 0 | 0 | 0 | 0 | 0 | 0 |
| K04 | 0 | 0 | 0 | 13638 | 0 | 0 | 1009 | 0 | 0 | 0 | 0 | 0 | 0 | 0 | 0 |
| K05 | 0 | 0 | 0 | 27362 | 0 | 62 | 0 | 0 | 0 | 0 | 0 | 0 | 0 | 0 | 0 |
| K06 | 0 | 0 | 0 | 19643 | 0 | 269 | 273 | 220 | 0 | 0 | 12 | 0 | 0 | 0 | 0 |
| K07 | 0 | 0 | 0 | 612 | 0 | 0 | 423 | 0 | 0 | 0 | 0 | 0 | 0 | 0 | 0 |
| K08 | 0 | 0 | 0 | 4503 | 0 | 0 | 0 | 0 | 0 | 0 | 0 | 0 | 0 | 0 | 0 |
| K09 | 0 | 0 | 0 | 6460 | 0 | 11 | 566 | 0 | 0 | 0 | 0 | 0 | 0 | 0 | 0 |
| K10 | 0 | 0 | 0 | 3663 | 0 | 10 | 0 | 0 | 0 | 0 | 0 | 0 | 0 | 0 | 0 |
| K11 | 0 | 0 | 0 | 1679 | 0 | 0 | 0 | 0 | 0 | 0 | 0 | 0 | 0 | 0 | 0 |
| K12 | 0 | 0 | 0 | 13738 | 45 | 1895 | 0 | 0 | 0 | 0 | 0 | 0 | 0 | 0 | 0 |
| K13 | 0 | 0 | 0 | 1249 | 104 | 305 | 0 | 464 | 0 | 0 | 0 | 0 | 43 | 0 | 18 |
| K14 | 0 | 0 | 0 | 815 | 0 | 700 | 0 | 0 | 0 | 0 | 0 | 0 | 0 | 0 | 0 |
| K15 | 0 | 0 | 0 | 32792 | 0 | 5525 | 0 | 0 | 0 | 0 | 0 | 0 | 0 | 0 | 0 |
| K16 | 0 | 0 | 0 | 1726 | 119 | 72 | 0 | 0 | 0 | 0 | 0 | 0 | 0 | 0 | 0 |
| K17 | 0 | 0 | 0 | 27974 | 0 | 398 | 344 | 337 | 0 | 0 | 0 | 0 | 0 | 0 | 0 |
| K18 | 0 | 0 | 0 | 6317 | 13 | 218 | 228 | 83 | 0 | 0 | 0 | 0 | 43 | 0 | 23 |
| K19 | 423 | 0 | 0 | 1042 | 107 | 424 | 0 | 1330 | 0 | 0 | 83 | 0 | 64 | 0 | 35 |
| K20 | 0 | 0 | 0 | 626 | 35 | 0 | 0 | 34 | 0 | 0 | 0 | 0 | 0 | 0 | 28 |
| K21 | 0 | 0 | 0 | 22561 | 269 | 249 | 0 | 1393 | 0 | 0 | 0 | 0 | 1917 | 0 | 0 |

This table shows eDNA metabarcoding result after bioinformatics processing. The number shows obtained reads. The “0” indicates no detection.

Table S5. Result of physical surveys.

| Site ID | *Cynops pyrrhogaster* | *Fejervarya limnocharis* | *Glandirana rugosa* | *Dryophytes japonica* | *Lithobates catesbeianus* | *Pelophylax nigromaculatus* | *P. porosus porosus* | *Rana japonica* |
| --- | --- | --- | --- | --- | --- | --- | --- | --- |
| A01 | 0 | 0 | 0 | 8 | 0 | 0 | 20 | 0 |
| A02 | 0 | 0 | 1 | 58 | 0 | 0 | 126 | 17 |
| A03 | 0 | 0 | 0 | 6 | 0 | 0 | 3 | 0 |
| A04 | 0 | 0 | 0 | 51 | 0 | 0 | 170 | 5 |
| A05 | 0 | 0 | 1 | 100 | 0 | 0 | 3 | 50 |
| A06 | 0 | 0 | 0 | 0 | 0 | 0 | 9 | 1 |
| A07 | 0 | 0 | 0 | 170 | 0 | 0 | 170 | 25 |
| A08 | 0 | 0 | 0 | 4 | 0 | 0 | 0 | 0 |
| A09 | 0 | 0 | 2 | 13 | 0 | 0 | 9 | 3 |
| A10 | 0 | 0 | 15 | 100 | 0 | 0 | 30 | 20 |
| A11 | NA | NA | NA | NA | NA | NA | NA | NA |
| A12 | NA | NA | NA | NA | NA | NA | NA | NA |
| A13 | NA | NA | NA | NA | NA | NA | NA | NA |
| A14 | NA | NA | NA | NA | NA | NA | NA | NA |
| B01 | NA | NA | NA | NA | NA | NA | NA | NA |
| B02 | 0 | 0 | 0 | 0 | 0 | 0 | 0 | 0 |
| B03 | 0 | 0 | 0 | 0 | 0 | 0 | 0 | 0 |
| B04 | NA | NA | NA | NA | NA | NA | NA | NA |
| B05 | NA | NA | NA | NA | NA | NA | NA | NA |
| B06 | 0 | 0 | 0 | 0 | 0 | 0 | 0 | 0 |
| B07 | 0 | 0 | 0 | 0 | 0 | 0 | 0 | 0 |
| B08 | NA | NA | NA | NA | NA | NA | NA | NA |
| B09 | NA | NA | NA | NA | NA | NA | NA | NA |
| B10 | 0 | 0 | 0 | 0 | 0 | 0 | 0 | 0 |
| B11 | 0 | 0 | 5 | 0 | 0 | 0 | 0 | 0 |
| C01 | NA | NA | NA | NA | NA | NA | NA | NA |
| C02 | NA | NA | NA | NA | NA | NA | NA | NA |
| C03 | NA | NA | NA | NA | NA | NA | NA | NA |
| C04 | NA | NA | NA | NA | NA | NA | NA | NA |
| C05 | 0 | 0 | 0 | 0 | 0 | 0 | 0 | 0 |
| C06 | 0 | 0 | 0 | 9 | 0 | 0 | 0 | 0 |
| C07 | 0 | 0 | 0 | 6 | 1 | 0 | 0 | 0 |
| C08 | 0 | 0 | 0 | 2 | 0 | 0 | 0 | 0 |
| C09 | 0 | 0 | 0 | 0 | 0 | 0 | 0 | 0 |
| C10 | 0 | 0 | 0 | 5 | 0 | 0 | 0 | 0 |
| C11 | NA | NA | NA | NA | NA | NA | NA | NA |
| C12 | 0 | 0 | 0 | 18 | 0 | 0 | 1 | 0 |
| C13 | 0 | 0 | 0 | 68 | 0 | 0 | 0 | 0 |
| C14 | 0 | 0 | 0 | 96 | 0 | 0 | 0 | 0 |
| C15 | 0 | 0 | 0 | 0 | 0 | 0 | 0 | 0 |
| C16 | 0 | 0 | 0 | 0 | 0 | 0 | 0 | 0 |
| C17 | 0 | 0 | 0 | 1 | 0 | 0 | 0 | 0 |
| C18 | 0 | 0 | 0 | 0 | 0 | 0 | 0 | 0 |
| C19 | 0 | 0 | 0 | 0 | 0 | 0 | 0 | 0 |
| C20 | NA | NA | NA | NA | NA | NA | NA | NA |
| D01 | NA | NA | NA | NA | NA | NA | NA | NA |
| D02 | NA | NA | NA | NA | NA | NA | NA | NA |
| D03 | 0 | 0 | 1 | 52 | 0 | 35 | 0 | 0 |
| D04 | 0 | 0 | 1 | 4 | 0 | 0 | 0 | 0 |
| D05 | 0 | 0 | 0 | 0 | 0 | 0 | 0 | 0 |
| D06 | 0 | 0 | 58 | 76 | 0 | 34 | 0 | 0 |
| D07 | 0 | 0 | 5 | 4 | 0 | 53 | 0 | 0 |
| D08 | 0 | 0 | 0 | 0 | 0 | 1 | 0 | 0 |
| D09 | NA | NA | NA | NA | NA | NA | NA | NA |
| D10 | NA | NA | NA | NA | NA | NA | NA | NA |
| D11 | NA | NA | NA | NA | NA | NA | NA | NA |
| D12 | NA | NA | NA | NA | NA | NA | NA | NA |
| D13 | 0 | 0 | 3 | 10 | 0 | 3 | 0 | 0 |
| D14 | 0 | 0 | 0 | 0 | 0 | 0 | 0 | 0 |
| D15 | 0 | 0 | 0 | 0 | 0 | 1 | 0 | 0 |
| D16 | 0 | 0 | 0 | 0 | 0 | 0 | 0 | 0 |
| D17 | 0 | 0 | 0 | 1 | 0 | 0 | 0 | 0 |
| D18 | 0 | 0 | 0 | 0 | 0 | 0 | 0 | 0 |
| D19 | NA | NA | NA | NA | NA | NA | NA | NA |
| D20 | NA | NA | NA | NA | NA | NA | NA | NA |
| F02 | 0 | 0 | 0 | 0 | 0 | 0 | 0 | 0 |
| F03 | 0 | 0 | 0 | 0 | 0 | 0 | 0 | 0 |
| F04 | 0 | 1 | 0 | 0 | 0 | 0 | 0 | 0 |
| F05 | 0 | 1 | 0 | 0 | 0 | 2 | 0 | 0 |
| F10 | 0 | 0 | 0 | 0 | 0 | 0 | 0 | 0 |
| F11 | 0 | 0 | 0 | 0 | 0 | 0 | 0 | 0 |
| F12 | 0 | 0 | 0 | 0 | 0 | 0 | 0 | 0 |
| F18 | 0 | 0 | 0 | 0 | 0 | 0 | 0 | 0 |
| F27 | 0 | 0 | 0 | 0 | 0 | 0 | 0 | 0 |
| F29 | 0 | 0 | 0 | 0 | 0 | 0 | 0 | 0 |
| F33 | NA | NA | NA | NA | NA | NA | NA | NA |
| F34 | NA | NA | NA | NA | NA | NA | NA | NA |
| G01 | 0 | 0 | 0 | 0 | 0 | 5 | 0 | 0 |
| G02 | 0 | 0 | 0 | 0 | 0 | 0 | 0 | 0 |
| G03 | 0 | 0 | 0 | 0 | 0 | 0 | 0 | 0 |
| G04 | 0 | 0 | 0 | 0 | 0 | 0 | 0 | 0 |
| G06 | 0 | 0 | 0 | 0 | 0 | 10 | 0 | 0 |
| G07 | 0 | 3 | 0 | 0 | 0 | 7 | 0 | 0 |
| G09 | 0 | 0 | 0 | 0 | 0 | 0 | 0 | 0 |
| G10 | 0 | 1 | 0 | 0 | 0 | 2 | 0 | 0 |
| G12 | 0 | 0 | 0 | 0 | 0 | 1 | 0 | 0 |
| G14 | 0 | 0 | 0 | 0 | 2 | 0 | 0 | 0 |
| G15 | 0 | 1 | 0 | 0 | 0 | 1 | 0 | 0 |
| G16 | 0 | 0 | 0 | 0 | 0 | 0 | 0 | 0 |
| G17 | 0 | 0 | 0 | 0 | 0 | 0 | 0 | 0 |
| G19 | 0 | 0 | 0 | 0 | 0 | 0 | 0 | 0 |
| G21 | 0 | 0 | 0 | 0 | 0 | 1 | 0 | 0 |
| G22 | 0 | 12 | 0 | 0 | 0 | 37 | 0 | 0 |
| G25 | 0 | 0 | 0 | 0 | 0 | 1 | 0 | 0 |
| G26 | 0 | 0 | 0 | 0 | 0 | 3 | 0 | 0 |
| G28 | 0 | 0 | 0 | 0 | 0 | 0 | 0 | 0 |
| G29 | NA | NA | NA | NA | NA | NA | NA | NA |
| H01 | 0 | 0 | 0 | 0 | 0 | 0 | 0 | 0 |
| H02 | 0 | 0 | 0 | 0 | 0 | 0 | 0 | 0 |
| H03 | 0 | 0 | 0 | 0 | 0 | 0 | 0 | 0 |
| H04 | 0 | 0 | 0 | 1 | 0 | 0 | 0 | 0 |
| H05 | 0 | 0 | 0 | 2 | 0 | 0 | 0 | 0 |
| H06 | 0 | 0 | 0 | 2 | 0 | 1 | 0 | 0 |
| H07 | NA | NA | NA | NA | NA | NA | NA | NA |
| H08 | NA | NA | NA | NA | NA | NA | NA | NA |
| H09 | NA | NA | NA | NA | NA | NA | NA | NA |
| H10 | NA | NA | NA | NA | NA | NA | NA | NA |
| I01 | 0 | 0 | 0 | 0 | 0 | 0 | 0 | 0 |
| I02 | 0 | 1 | 0 | 0 | 0 | 0 | 0 | 0 |
| I03 | 0 | 1 | 0 | 0 | 0 | 0 | 0 | 0 |
| I04 | 0 | 10 | 0 | 0 | 0 | 0 | 0 | 0 |
| I05 | 0 | 0 | 0 | 0 | 0 | 0 | 0 | 0 |
| I06 | 0 | 0 | 0 | 0 | 0 | 0 | 0 | 0 |
| I07 | 0 | 0 | 0 | 0 | 0 | 0 | 0 | 0 |
| I08 | 0 | 0 | 0 | 0 | 0 | 0 | 0 | 0 |
| I09 | NA | NA | NA | NA | NA | NA | NA | NA |
| I10 | NA | NA | NA | NA | NA | NA | NA | NA |
| I11 | NA | NA | NA | NA | NA | NA | NA | NA |
| I12 | NA | NA | NA | NA | NA | NA | NA | NA |
| J01 | 2 | 0 | 1 | 0 | 0 | 0 | 0 | 0 |
| J02 | NA | NA | NA | NA | NA | NA | NA | NA |
| J03 | 6 | 0 | 0 | 0 | 0 | 1 | 0 | 0 |
| J04 | NA | NA | NA | NA | NA | NA | NA | NA |
| J05 | NA | NA | NA | NA | NA | NA | NA | NA |
| J06 | NA | NA | NA | NA | NA | NA | NA | NA |
| J09 | 32 | 0 | 1 | 0 | 0 | 0 | 0 | 0 |
| J10 | 0 | 0 | 0 | 0 | 0 | 0 | 0 | 0 |
| J11 | 0 | 0 | 0 | 0 | 0 | 0 | 0 | 0 |
| J12 | 1 | 0 | 0 | 0 | 0 | 0 | 0 | 0 |
| J13 | 4 | 0 | 0 | 0 | 0 | 1 | 0 | 0 |
| J14 | 5 | 0 | 0 | 0 | 0 | 0 | 0 | 1 |
| J19 | 23 | 0 | 1 | 1 | 0 | 0 | 0 | 0 |
| J20 | 23 | 0 | 0 | 0 | 0 | 1 | 0 | 0 |
| J21 | 0 | 0 | 0 | 0 | 0 | 0 | 0 | 0 |
| J22 | 0 | 0 | 1 | 0 | 0 | 0 | 0 | 1 |
| J23 | 12 | 0 | 1 | 0 | 0 | 0 | 0 | 0 |
| J24 | 15 | 0 | 0 | 0 | 0 | 0 | 0 | 0 |
| J25 | 0 | 0 | 0 | 0 | 0 | 0 | 0 | 0 |
| J32 | 6 | 0 | 2 | 0 | 0 | 0 | 0 | 1 |
| K01 | 0 | 0 | 0 | 0 | 0 | 0 | 0 | 0 |
| K02 | 0 | 0 | 0 | 0 | 0 | 0 | 0 | 0 |
| K03 | 0 | 0 | 0 | 0 | 0 | 0 | 0 | 0 |
| K04 | 0 | 0 | 0 | 0 | 0 | 0 | 0 | 0 |
| K05 | 0 | 0 | 0 | 0 | 0 | 0 | 0 | 0 |
| K06 | 0 | 0 | 0 | 0 | 0 | 0 | 0 | 0 |
| K07 | 0 | 0 | 0 | 0 | 0 | 0 | 0 | 0 |
| K08 | 0 | 0 | 0 | 0 | 0 | 0 | 0 | 0 |
| K09 | 0 | 0 | 0 | 0 | 0 | 0 | 0 | 0 |
| K10 | 0 | 0 | 0 | 0 | 0 | 0 | 0 | 0 |
| K11 | 0 | 0 | 0 | 0 | 0 | 0 | 0 | 0 |
| K12 | 0 | 0 | 0 | 0 | 0 | 0 | 0 | 0 |
| K13 | 0 | 0 | 0 | 0 | 0 | 0 | 0 | 0 |
| K14 | 0 | 0 | 0 | 0 | 0 | 0 | 0 | 0 |
| K15 | 0 | 0 | 0 | 0 | 0 | 0 | 0 | 0 |
| K16 | 0 | 0 | 0 | 0 | 0 | 0 | 0 | 0 |
| K17 | 0 | 0 | 0 | 0 | 0 | 0 | 0 | 0 |
| K18 | 0 | 0 | 0 | 0 | 0 | 0 | 0 | 0 |
| K19 | 0 | 0 | 0 | 0 | 0 | 0 | 0 | 0 |
| K20 | 0 | 0 | 0 | 0 | 0 | 0 | 0 | 0 |
| K21 | 0 | 0 | 0 | 0 | 0 | 0 | 0 | 0 |

The number shows captured individuals. The “0” indicates no detection. The “NA” indicates no performing physical surveys.
